# Improved computational identification of drug response using optical measurements of human stem cell derived cardiomyocytes in microphysiological systems

**DOI:** 10.1101/787390

**Authors:** Karoline Horgmo Jæger, Verena Charwat, Bérénice Charrez, Henrik Finsberg, Mary M. Maleckar, Samuel Wall, Kevin E. Healy, Aslak Tveito

## Abstract

Cardiomyocytes derived from human induced pluripotent stem cells hold great potential for drug screening applications. However, their usefulness is limited by the relative immaturity of cells’ electrophysiological properties as compared to native cardiomyocytes in the adult human heart. In this work, we extend and improve on methodology to address this limitation, building on previously introduced computational procedures which predict drug effects for mature cells based on changes in optical measurements of action potentials and Ca^2+^ transients made in stem cell derived cardiac microtissues. This methodology quantifies ion channel changes through the inversion of data into a mathematical model, and maps this response to a mature phenotype through the assumption of functional invariance of fundamental intracellular and membrane channels during maturation.

Here we utilize an updated action potential model to represent both immature and mature cells, apply an IC50-based model of dose-dependent drug effects, and introduce a continuation-based optimization algorithm for analysis of dose escalation measurements using five drugs with known effects. The improved methodology can identify drug induced changes more efficiently, and quantitate important metrics such as IC50 in line with published values. Consequently, the updated methodology is a step towards employing computational procedures to elucidate drug effects in mature cardiomyocytes for new drugs using stem cell-derived experimental tissues.

## 1 Introduction

The development of human induced pluripotent stem cells (hiPSCs) opens promising avenues of investigation into a wide variety of fundamental questions in cell physiology and beyond (for recent reviews, see e.g., [1, 2, 3]). One of the more immediately tractable applications of hiPSCs is the creation of specific human cell and tissue samples to augment drug discovery and development pipelines. These pipelines have traditionally relied on animal models in key areas of testing, but is limited by significant physiological differences between animal and human cells (see e.g., [1, 3, 4, 5]). These differences, both at the genetic and proteomic levels, give rise to distinctly non-human system dynamics, for example, a mouse’s heart rate is much more rapid than a human’s (∼600 bpm vs. ∼60 bpm), such that it is often difficult to translate drug effects from one species to another (see e.g., [1, 3, 4, 5]).

By using hiPSC-derived cells, it is possible to measure drug effects directly in human-based systems, and therapeutics can eventually, in principle, be tested and adjusted at the level of the individual patient. This hiPSC-based, patient-centric approach opens up great possibilities for drug development, both in terms of the scope of illnesses approachable, including disorders caused by rare mutations, as well as improved safety by the early identification of drug side effects in *human* cells. Nevertheless, hiPSCs are also associated with a variety of scientific challenges that must be resolved to realize the full potential of the technology (see e.g., [4, 6, 7, 8, 9, 10]).

Maturity of generated cells and tissues is one of these key challenges, a prominent example being the maturation of hiPSC-derived cardiomyocytes (hiPSC-CMs) [11]. Human cardiomyocytes develop over many years (see [12], ch. 21), and during this period the density of specific ion channels changes significantly, due both to increased area of the cell membrane and proliferation of membrane channels (see e.g., [13, 14, 15]). The physiological response of immature hiPSC-CMs to a drug cannot necessarily be used to infer the properties of the drug, nor the response of adult human cardiomyocytes. Even if it is known exactly how a drug affects an immature hiPSC-CM, it is difficult to deduce its effect on adult cells; direct interpretation may in fact lead to both false positives and false negatives (see [10, 16]).

In [17], we used mathematical modeling of cardiac cell dynamics to address these challenges associated with the application of hiPSC-CMs. Such mathematical modeling of the cardiac action potential (AP) remains an active area of research, and sophisticated models have been developed in order to accurately simulate both single cells and cardiac tissue dynamics (see e.g., [18, 19, 20, 21, 22, 23, 24, 25, 26, 27]). We presented an algorithm for inverting experimental measurements of the membrane potential and the cytosolic calcium (Ca^2+^) concentration in order to obtain parameters for a mathematical model of the hiPSC-CM AP. We then demonstrated how this model of immature cells (hiPSC-CMs) can be mapped to an AP model representing mature cells. We were able to estimate the effect of a drug on essential ion currents for hiPSC-CMs as based on measurements from a microphysiological system [10], and then to map this effect onto the adult cardiomyocyte AP model. The combination of these two methods permitted, in principle, to deduce drug effects on mature cells (adult human cardiomyocytes) as based on measurements of immature cells (hiPSC-CMs in a microphysiological system). The overall method developed in [17] is illustrated in Figure 1. In this procedure, we take optical measurements using fluorescent dyes in a microphysiological system to define relative traces of the membrane potential and the cytosolic Ca^2+^ concentration for cells under normal media conditions and in the presence of drugs. We then define a mathematical model for the control (undrugged) cases by identifying parameters denoted by *p*^IM,c^ (IM is for immature, c is for control) in an AP model that matches the experimental waveforms. Using this model of immature cells, we then define a maturation matrix *Q* such that *Qp*^IM,c^ = *p*^M,B^, where *p*^M,B^ (M is for mature, B is for base) are *known* parameters representing a generic AP model of an adult human cardiomyocyte. Here, the matrix *Q* represents the developmental change in ion channel density and geometry from IM to M, independent of drug effects on individual channels.

**Figure 1:**
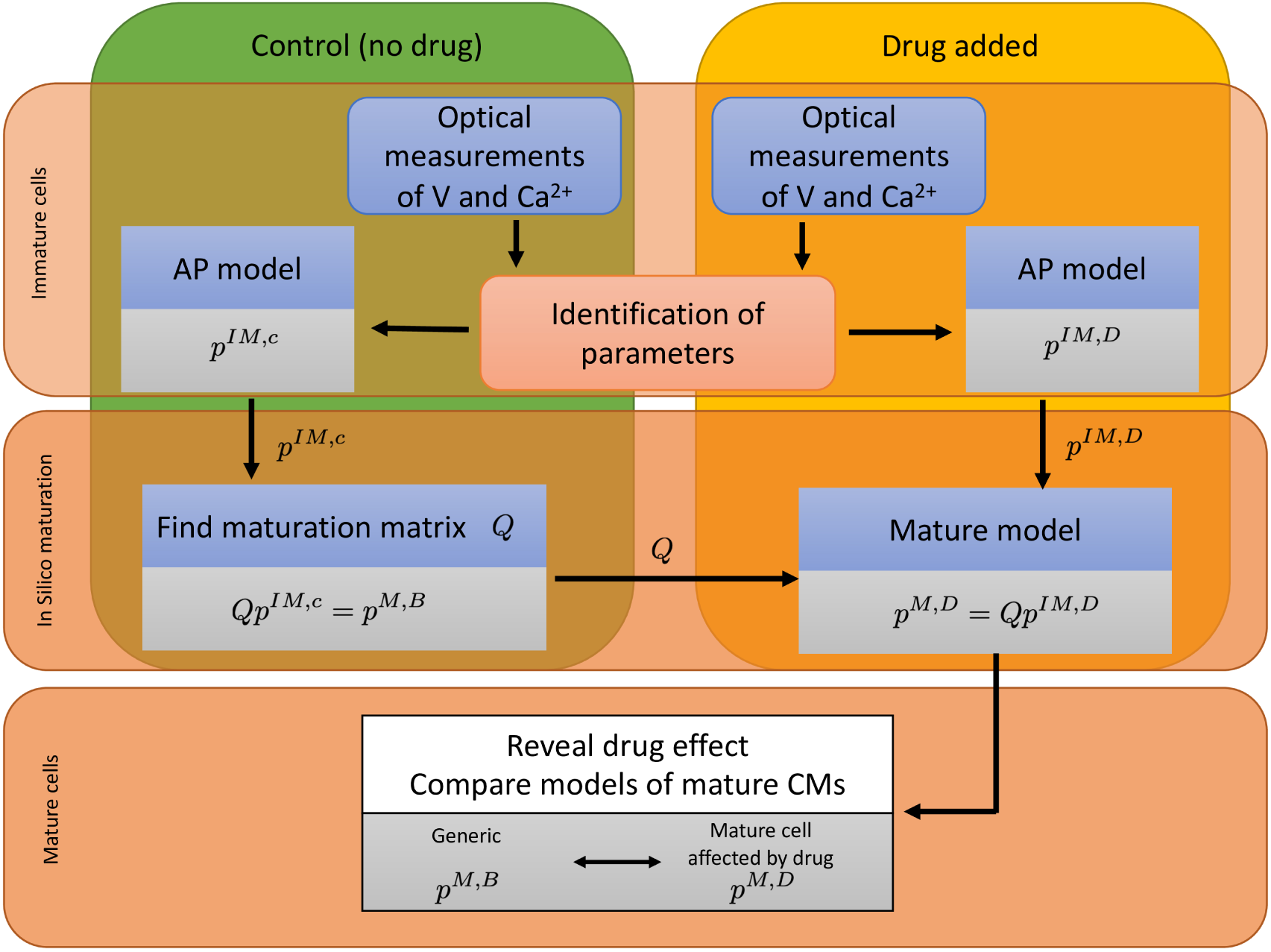
The effect of a drug on mature cardiomyocytes can be identified by the process illustrated. The cytosolic calcium concentration (Ca^2+^) and the membrane potential (V) are measured in a microphysiological system ([4, 10]) of immature cells (hiPSC-CMs). Data are collected when no drug has been applied (control, c) and when a drug has been applied (D). The data are used to parameterize a model for both cases, represented by the parameter vectors *p*^*IM,c*^ and *p*^*IM,D*^ for the control and drugged cases, respectively. The control parameter vector *p*^*IM;c*^ is used to define the maturation matrix *Q* such that *Q*^*pIM,c*^ = *p*^*M,B*^, where *p*^*M,B*^ is the parameter vector of a generic base model of mature, adult human cardiomyocytes. By comparing the mature parameter vectors for the control and drugged cases, the effect of the drug can be identified.

Next, experimental traces of the membrane potential and the cytosolic Ca^2+^ concentration are taken for the same cells in the presence of drugs, and these traces are used to define a new parameter-vector *p*^IM,D^ (Immature, Drug) that matches the data. This new parameterization gives us information about what modeled channels have been altered by the drug. Then, by assuming that the drug affects *every individual ion channel* in the same manner for the IM and M cells, the parameter vector for the mature case is given by *p*^M,D^ = *Qp*^IM,D^. *Hence, we can find an AP model for mature human cardiomyocytes under the influence of the drug, even though only the effect in the immature hiPSC-CM case has been measured.*

The present report aims to present a number of modifications to improve the accuracy and reliability of these methods. First, using the AP models of O’Hara et al. [18], Grandi et al. [19], and Paci et al. [28, 29, 30, 31] as a basis, we have derived a new AP model to improve representation of experimental data. As our inversion algorithm is based on conducting a huge number of simulations with varying parameter values, it is essential to have a model that is stable with respect to perturbations of the parameters. Therefore, the new model is designed for improved stability. In particular, the model of the intracellular Ca^2+^ dynamics has been modified to avoid instabilities in the balance between the influx and efflux of Ca^2+^ to the sarcoplasmic reticulum (SR).

In addition, our aim has been to create models that can be mapped back and forth between immature (hiPSC-CM) and mature (adult cardiomyocyte) cases. A vital modeling assumption has been that the individual channels are the same in the immature and mature cases, and that only channel density should change. However, existing AP models are not derived with such a mapping in mind, and models of identical single channel dynamics vary significantly among models. Therefore, we have derived a new AP model which strictly adheres to the principle that every current (and flux) should be written as a product of the ion channel density and the dynamics of a single channel; identical ion channels are represented by identical mathematical models. Consequently, the mathematical representation of *a single channel* is the same for the IM and M cases in the novel AP model presented here. Finally, we present a new method for inverting experimental data into parameters for the AP model by introducing a continuation-based approach, searching for optimal parameters by gradually moving from known parameters to the parameters we want to identify. Continuation methods are well developed in scientific computing (see e.g., [32, 33]) and offer significant computational savings to find optimal solutions.

In this manuscript, we first motivate and describe the approaches outlined above. We then evaluate these methods with respect to accuracy using simulated data. Subsequently, the new methods are used to identify the effect of five well-characterized drugs based on optical measurements of hiPSC-CMs. In all cases considered, the predicted effects are consistent with known drug effects, lending credence to the principle that novel drug effects on adult cardiomyocytes could reliably be estimated using measurements of hiPSC-CMs and the described methodology.

## 2 Methods

Here we offer a detailed presentation of all steps illustrated in Figure 1. First, we present the derivation of a new AP model. Next, we describe the inversion method used in our computations. Finally, we discuss how to characterize the identifiability of the parameters involved in the inversion as based on singular value decomposition (SVD) of model currents.

### 2.1 The base model

As noted above, we aim to define an AP model that can be scaled from very early stages of human development (days) to fully developed adult heart cells. To review, for one specific membrane current, we assume that the only difference between the immature (IM) case and the mature (M) case is that the number of channels and the membrane area has changed; thus, the density of the specific ion channel carrying the current has changed, but the properties of every individual channel remains the same. The same principle holds for the intracellular Ca^2+^ machinery; the individual channels and buffers remain the same, but both the intracellular volumes and the number of channels change from IM to M. Our model will, therefore, be based on models of single ion channel dynamics and *only the density of these single channels will change from IM to M*. When a drug is involved, we assume that the effect of the drug on a *single channel* is the same in the IM and M cases, and therefore one can use the effect in the IM case to estimate the effect for the M case.

The membrane currents and intracellular compartments of the base model are illustrated in Figure 2. In the formulation of the base model, the membrane potential (*v*) is given in units of mV, and the Ca^2+^ concentrations are given in units of mM. All currents are expressed in units of A/F, and the Ca^2+^ fluxes are expressed as mmol/ms per total cell volume (i.e., in units of mM/ms). Time is given in ms.

**Figure 2:**
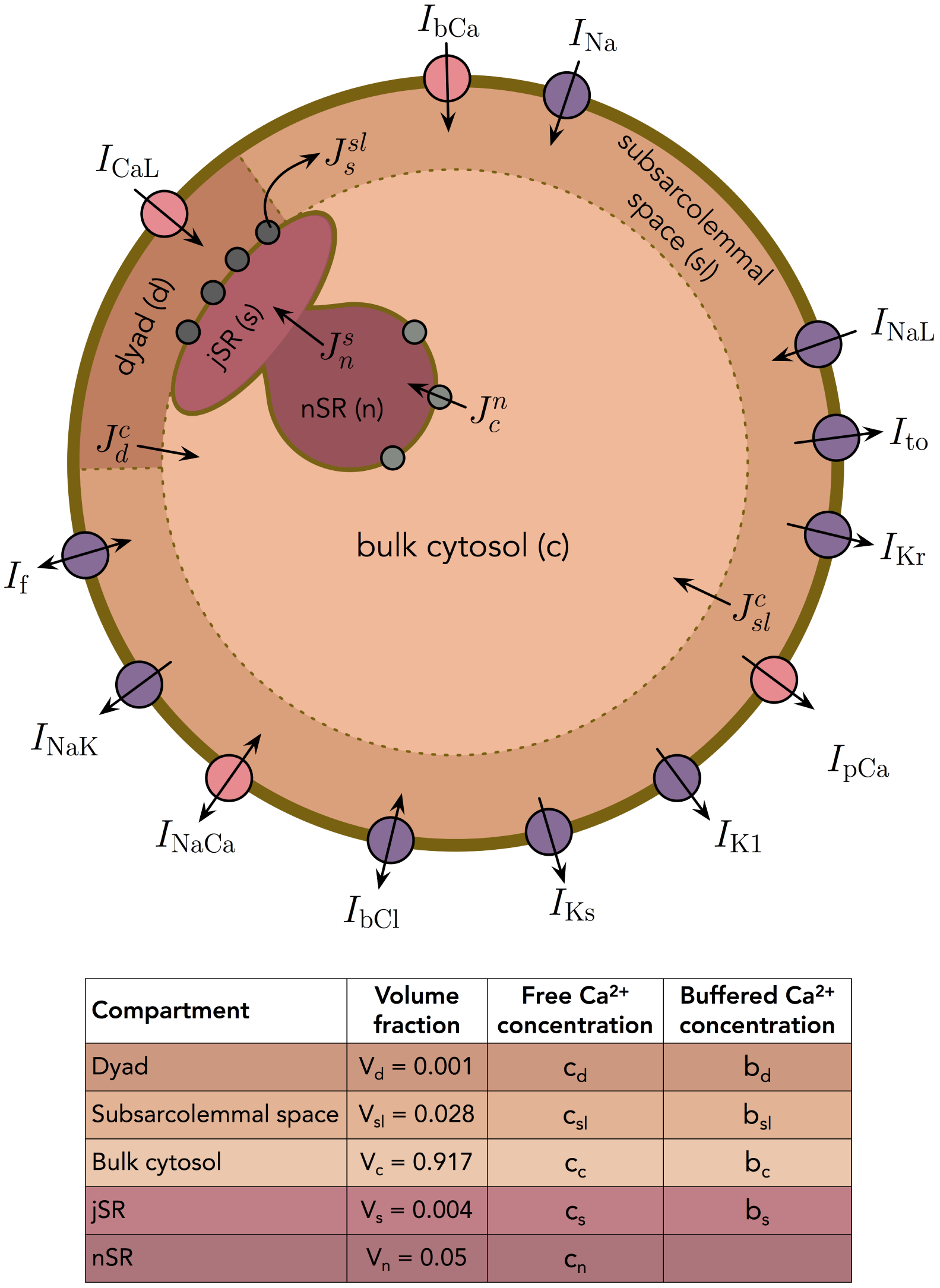
Membrane currents, Ca^2+^ fluxes and intracellular compartments of the base model.

### 2.2 Modeling the membrane currents

The standard model (see e.g., [34, 35, 36, 37]) of the membrane potential of an excitable cell is given by the equation

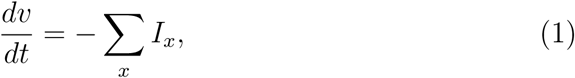

where *v* is the membrane potential (in mV), and *I*_*x*_ are the membrane currents through ion channels of different types, as well as pumps and exchangers located on the cell membrane.

These currents are all given in units of A/F, and may be written on the form

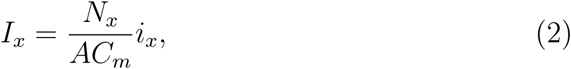

where *N*_*x*_ is the number of channels of type *x* on the cell membrane, *A* is the area of the cell membrane (in *µ*m^2^) and *C*_*m*_ is the specific capacitance of the cell membrane (in pF/*µ*m^2^). Furthermore, *i*_*x*_ represents the average current through a single channel of type *x* (in pA). For voltage-gated ion channels, this average single-channel current is given on the form

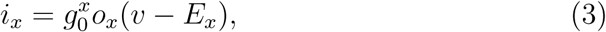

where 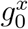 is the conductance of a single open channel (in nS), *E*_*x*_ is the equilibrium potential of the channel (in mV), and *o*_*x*_ is the unitless open probability of the channel. Note that in models given on this form, it is common to consider a lumped parameter *g*_*x*_, given by

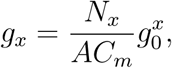

and parameters of this type are given for each of the ion channels considered in the base model in the Supplementary information. For membrane pumps and exchangers, the single-channel current is given on a similar form. The specific currents included in the model will be described below.

#### 2.2.1 Scaling of the membrane currents

As mentioned above, we assume that the specific membrane capacitance and the ion channels responsible for each of the membrane currents are the same during different stages of development for the cell, but that the number of ion channels, *N*_*x*_, and the membrane area, *A*, may differ. Therefore, currents can be mapped from one stage of development, *S*_1_, to another stage of development, *S*_2_, simply by adjusting the channel density of the currents. More specifically, for the formulation (1)–(2), this means that we assume that the parameter *C*_*m*_ and the expressions for the single-channel currents *i*_*x*_, are the same for *S*_1_ and *S*_2_, but that the channel density 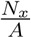, can be different. Let 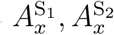 and 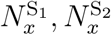 denote the membrane area and number of channels of type *x* for the *S*_1_ and *S*_2_ cases, respectively. Furthermore, we let *λ*_*x*_ represent the change of channel density in the sense that

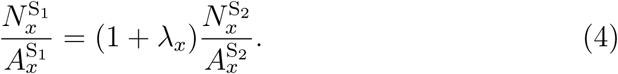

Now, the *S*_1_ and *S*_2_ currents are related according to

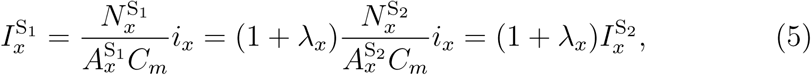

for each of the currents *x*.

#### 2.2.2 The base model is the generic mature model

Based on these considerations, it is convenient to define *one* default base model from which all other models are derived to simplify a mapping procedure between different development stages.

Defining a base model as representing IM cells, from which M cells subsequently develop, may seem to be a natural choice. However, in the scheme illustrated in Figure 1, there is only one fixed model – the generic M model – while all other models will change depending on the experimental measurements. For simplicity, we therefore define the generic M model to be the *default base model*, and scale all other models relative to this model.

#### 2.2.3 Main currents present in human cardiomyocytes

Modern models of human cardiomyocytes are complex and the models for the individual currents are based on years of experience using patch-clamp measurements. In the formulation (1), our aim has been to include the main currents present in human cardiomyocytes, but to keep the number of currents as low as feasible in order to keep the base model relatively simple. The experimental inputs in the present methodology are optically-derived, and data based on sensitive dyes are not expected to be able to uncover equally fine details of the dynamics as compared to traditional electrophysiological measurements derived via patch clamp. It is therefore reasonable to represent the data using simpler models. Our choice of currents is based on the O’Hara-Rudy et al. model [18] and the Grandi et al. model [19] for human mature ventricular cardiomyocytes, in addition to the Paci et al. model [28] for hiPSC-CMs. Furthermore, we have focused on including currents considered to be critical for depolarization and repolarization of the AP and, therefore, those typically investigated for response to drugs (see e.g., [38]).

In [38], the fast sodium current, *I*_Na_, the late sodium current, *I*_NaL_, the L-type Ca^2+^ current, *I*_CaL_, the transient outward potassium current, *I*_to_, the rapid and slow delayed rectifier potassium currents, *I*_Kr_ and *I*_Ks_, and the inward rectifier potassium current, *I*_K1_, were investigated for their drug responses, and we have included each of these currents in our model. In addition, we have included the sodium-potassium pump, *I*_NaK_, the sodium-calcium exchanger, *I*_NaCa_, the Ca^2+^ pump, *I*_pCa_, the background Ca^2+^ current, *I*_bCa_, and the background chloride current, *I*_bCl_, as they all appear to have a significant effect on the computed AP and Ca^2+^ transient of the Grandi et al. model [19]. Furthermore, we have included the hyperpolarization-activated cyclic nucleotide-gated funny current, *I*_f_. While this current is very small for mature ventricular cardiomyocytes, it is substantial for hiPSC-CMs [39]. The formulation used for each of the currents is given in the Supplementary information. The formulations are based on those of the currents in the Paci et al. model [28], the Grandi et al. model [19], and the O’Hara-Rudy et al. model [18].

### 2.3 Modeling intracellular Ca^2+^ dynamics

In addition to the membrane potential, we also want the base model to represent the dynamics of the intracellular Ca^2+^ concentration. We consider the following five intracellular compartments:

1. The dyad, representing the small cytosolic subspace between the L-type Ca^2+^ channels and the ryanodine receptors (RyRs),
2. The subsarcolemmal space, representing the remaining part of the cytosolic space that is located close to the membrane,
3. The bulk cytosolic space,
4. The junctional sarcoplasmic reticulum (jSR), representing the part of the SR that is close to the RyR-channels,
5. The network sarcoplasmic reticulum (nSR), representing the remaining part of the SR.

The Ca^2+^ concentrations and volume fractions defined for each of these compartments are given in Figure 2. In all compartments except the nSR, we consider both the concentration of free Ca^2+^ and the concentration of Ca^2+^ bound to a buffer. The Ca^2+^ concentration in the extracellular space is assumed to remain constant. The intracellular Ca^2+^ fluxes between compartments are illustrated in Figure 2 (and membrane currents involving exchange of Ca^2+^-ions between the intracellular and extracellular spaces are marked with pink circles). All the Ca^2+^ fluxes considered in the base model are summarized in Table 1.

**Table 1:** Ca^2+^ fluxes of the base model. The direction of all membrane fluxes are defined such that a positive flux corresponds to Ca^2+^ ions flowing into the cell.

| Flux | Description |
| --- | --- |
| $J_s^{sl}$ | Flux through the RyRs from the jSR to the SL |
| $J_c^n$ | Flux through the SERCA pumps from the bulk cytosol to the nSR |
| $J_d^c$ | Passive diffusion flux between the dyad and the bulk cytosol |
| $J_{sl}^c$ | Passive diffusion flux between the SL and the bulk cytosol |
| $J_n^s$ | Passive diffusion flux between the nSR and the jSR |
| $J_d^b$ | Free $\text{Ca}^{2+}$ binding to a buffer in the dyad |
| $J_{sl}^b$ | Free $\text{Ca}^{2+}$ binding to a buffer in the SL |
| $J_c^b$ | Free $\text{Ca}^{2+}$ binding to a buffer in the bulk cytosol |
| $J_s^b$ | Free $\text{Ca}^{2+}$ binding to a buffer in the jSR |
| $J_{\text{CaL}}$ | $\text{Ca}^{2+}$-flux through the L-type $\text{Ca}^{2+}$ channels from the extracellular space to the dyad |
| $J_{\text{bCa}}$ | Background $\text{Ca}^{2+}$ flux from the extracellular space to the SL |
| $J_{\text{pCa}}$ | $\text{Ca}^{2+}$ flux through the $\text{Ca}^{2+}$ pump between the extracellular space and the SL |
| $J_{\text{NaCa}}$ | $\text{Ca}^{2+}$ flux through the $\text{Na}^+$-$\text{Ca}^{2+}$ exchanger between the extracellular space and the SL |
| $J_e^{sl}$ | Total $\text{Ca}^{2+}$ flux from the extracellular space to the SL, defined as $J_e^{sl} = J_{\text{bCa}} + J_{\text{pCa}} + J_{\text{NaCa}}$. |

#### 2.3.1 Modeling release from the SR

In our model of Ca^2+^ dynamics, we deviate from previous modeling approaches in two specific ways:

1. Ca^2+^ is released through RyR channels from the SR directly to the subsarcolemmal space (SL) and not to the dyad.
2. Release of Ca^2+^ through the RyR channels is a product of two factors; one factor models the open probability of the RyR channels, whereas the other models the availability of channels that can be opened. We assume that each channel can only process a certain amount of Ca^2+^ before it deactivates.

We will see below that these two modeling assumptions lead to a model that exhibits two key physiological features of Ca^2+^ release from the SR of cardiomyocytes, so-called *high gain* and *graded release* (see Section 4.2.2 for explanations of these terms).

#### 2.3.2 Definition of Ca^2+^ fluxes

As noted above, all Ca^2+^ fluxes, *J*, are defined in terms of the number of ions flowing per time per total cell volume, in units of mM/ms. Accordingly, the size of a flux in mmol/ms is given by 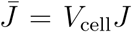, where *V*_cell_ is the cell volume (in L).

Similarly, for a single compartment with volume 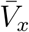 (in L), volume fraction (dimensionless) 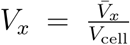 and Ca^2+^ concentration *c*_*x*_ (in mM), the total number of Ca^2+^ ions in the compartment (in mmol) is given by 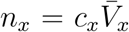. The change in the number of Ca^2+^-ions in the compartment is given by

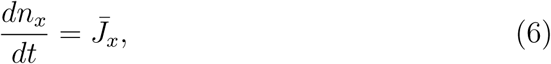

where 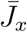 is the flux of ions into the compartment given in mmol/ms. It is also useful to define an associated concentration flux per total cell volume, 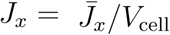 (in mM/ms). Dividing both sides of (6) by the compartment volume 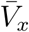, we obtain the following equation for the change in Ca^2+^ concentration:

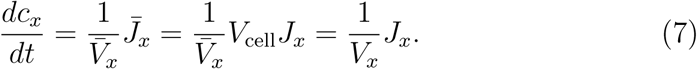

Expanding this approach to all the compartments and fluxes of the model, we obtain the following system of equations for the intracellular Ca^2+^ dynamics:

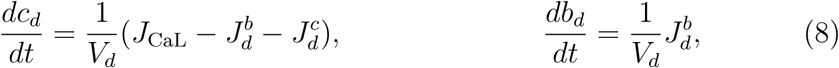

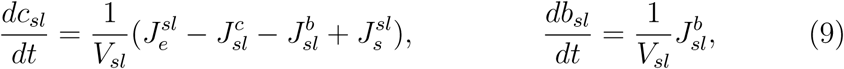

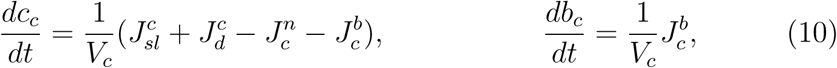

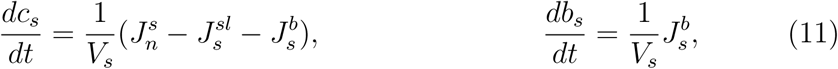

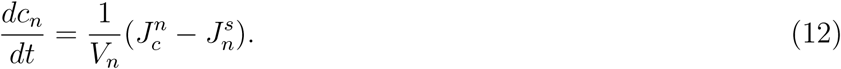

The expressions for each of the Ca^2+^ fluxes are defined below.

##### Expressions for fluxes through ion channels

Every flux *J* = *J*_*x*_ in the model representing fluxes through a type of channel will be written on the form

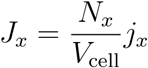

where *N*_*x*_ is the number of channels of type *x*, and *j*_*x*_ is the average flux through a single channel of type *x*.

##### Flux through the SERCA pumps 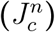

The flux from the bulk cytosol to the nSR through SERCA is given on the form

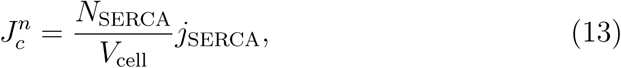

where *N*_SERCA_ is the number of SERCA pumps on the membrane of the nSR, *V*_cell_ is the total cell volume (in L) and *j*_SERCA_ is the flux through a single SERCA pump (in mmol/ms). The flux through a single pump is given by an expression based on the formulation in the Grandi et al. model [19]:

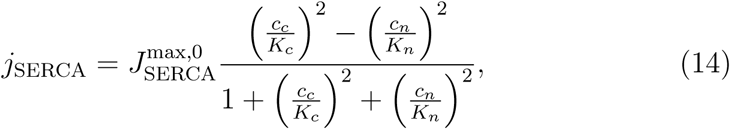

where 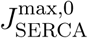 has unit mmol/ms and *K*_*c*_ and *K*_*n*_ have unit mM. Defining the parameter

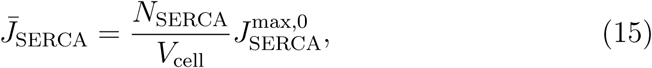

with unit mM/ms, the SERCA flux may be written as

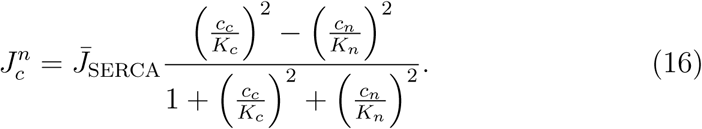

##### Scaling the SERCA flux

Like for the maturation process with respect to sarcolemmal ion channels, we also assume that cells of different levels of maturity may have different geometries and different densities of SERCA pumps, but that the function of each individual SERCA pump is the same. This means that we assume that the expression for the flux through a single SERCA pump, *j*_SERCA_, remains the same, but that the factor 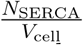 may differ between immature and mature cells. We again represent the change in the SERCA pump density by introducing a scaling factor *λ*_SERCA_ between one stage of maturity, *S*_1_, to another stage, *S*_2_, such that

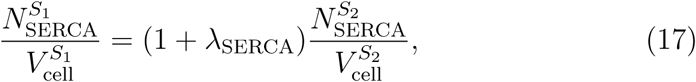

Where

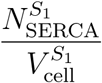

is the SERCA pump density in the model for maturity stage *S*_1_ and

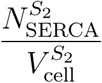

is the density in the model for maturity stage *S*_2_. In the model formulation, this can be represented on the form

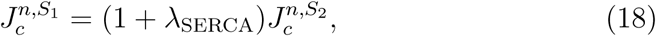

where 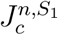 is the expression for the SERCA pump flux in the *S*_1_ state and 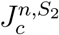 is the expression in the *S*_2_ state.

##### Flux through RyRs 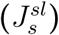

As we will employ the base model for several different parameter combinations, the model for the RyR flux must be stable, in the sense that careful tuning of the model is not requisite to ensure reasonable activation and deactivation of the RyRs.

As outlined above, we let the Ca^2+^ released from the SR enter the SL space rather than the dyad. This is done in order to achieve graded release (see Section 4.2.2), in the sense that the amount of Ca^2+^ released from the SR through the RyRs should depend directly upon the amount of Ca^2+^ entering the cell through L-type Ca^2+^ channels. If the model were to be formulated such that Ca^2+^ released from the jSR instead entered the dyad, it would be difficult to distinguish the increase in dyadic Ca^2+^ concentration resulting from L-type Ca^2+^ channel flux as opposed to release via RyRs. Directing the RyR flux into the SL, the concentration change in the dyad is almost exclusively due to the influx through L-type Ca^2+^ channels, and by letting the flux through the RyRs depend on the Ca^2+^ concentration in the dyad, we achieve graded release.

Furthermore, a common modeling approach for the RyRs is to govern inactivation by a decreased jSR concentration (see e.g., [40]). However, for large variations in parameter values, this may lead to model instabilities, because the jSR concentration depends upon the balance between the flux through the SERCA pumps and the RyRs, which depend upon the balance between the Ca^2+^ fluxes in and out of the cell. In order to avoid an RyR model whose inactivation mechanism depends on the jSR concentration, we instead introduce a new assumption that some RyRs are only able to carry a given amount of Ca^2+^ ions during each AP.

We then assume that a small portion of the RyR channels are always open (type 0), while the remaining channels (type 1) are activated by an increased dyadic Ca^2+^ concentration and are inactivated after they have transported a given amount of Ca^2+^ ions. Therefore, the total flux through the RyRs may be expressed as

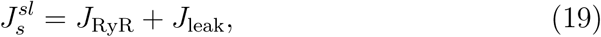

where *J*_RyR_ represents the flux through the RyR channels of type 1 and *J*_leak_ represents the flux through the RyR channels of type 0. We assume that the flux through the two types of RyR channels are given by expressions of the form

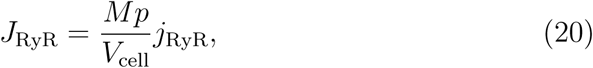

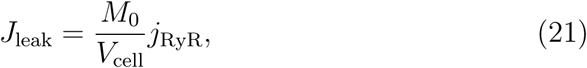

where *j*_RyR_ denotes the flux through a single open RyR channel (in mmol/ms) and *V*_cell_ denotes the total cell volume (in L). In addition, *M*_0_ denotes the number of RyR channels that are always open (type 0), *M* denotes the number of available RyR channels of type 1, and *p* is the open probability of the channels of type 1. The single channel flux through the RyRs is given by

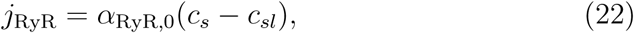

where *α*_RyR,0_ (in L/ms) represents the rate of release. Furthermore, the open probability of the RyR channels of type 1 is modeled by a simple function that increases sigmoidally with the dyadic Ca^2+^ concentration, *c*_*d*_, based on the model in [41]:

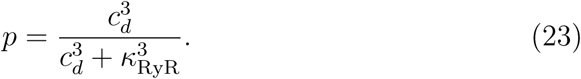

We let the total number of RyR channels of type 1 be given by *N*_RyR_ and the total number of RyR channels of type 0 be given by

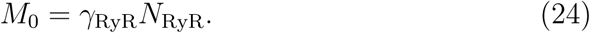

In other words, the total number of RyR channels (of both types) is given by (1 + *γ*_RyR_)*N*_RyR_.

We assume that every RyR of type 1 is able to transport a fixed amount *Z* Ca^2+^ ions during an AP. After Z ions have been transported, the channel becomes inactivated. However, we assume that as the dyadic Ca^2+^ concentration, *c*_*d*_, returns to rest and the open probability, *p*, consequently decreases, the inactivated channels gradually recover from inactivation. We let the number of available channels of type 1 be governed by

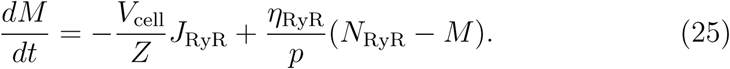

Here, the first term dominates for large values of *p*, driving *M* towards zero as more Ca^2+^ is transported through the RyR channels of type 1. Furthermore, for small values of *p* (i.e., at rest), the second term dominates and drives *M* towards the maximum value *N*_RyR_.

In order to reduce the number of free parameters in the model, we define a scaled variable *r*, defined as 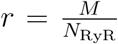, and divide both sides of equation (25) by *N*_RyR_. The equation then reads

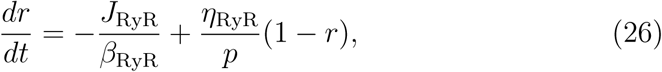

Where

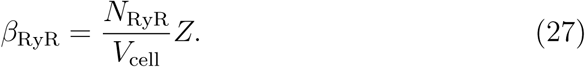

Inserting *M* = *rN*_RyR_ into (20) and defining

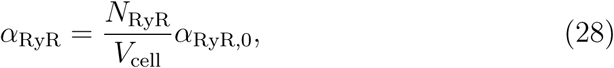

we get the following expression for active RyR flux

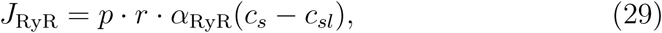

where we recall that

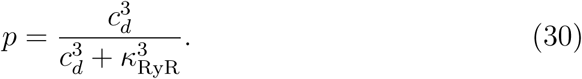

Moreover, inserting (24) and (28) into (21), we obtain

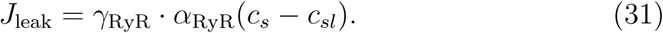

##### Scaling the RyR flux

When considering cells of different levels of maturity, we again assume that the number of RyRs and the cell volume may be different, but that the function of a single RyR channel is the same for different levels of maturity. We also assume that the ratio between RyR channels of type 0 and 1, *γ*_RyR_, and the number of Ca^2+^ ions that each RyR channel of type 1 can transport, *Z*, is the same for the different maturity levels. Considering the model (26)–(31), this means that the only adjustment necessary between two maturity levels *S*_1_ and *S*_2_ is an adjustment of the density 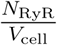 in the definition of *α*_RyR_ and *β*_RyR_. We therefore introduce a scaling factor *λ*_RyR_ such that

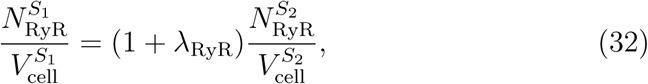

and represent this adjustment of the RyR density in the model by scaling *α*_RyR_ and *β*_RyR_ by

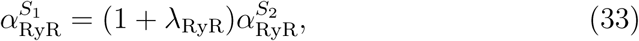

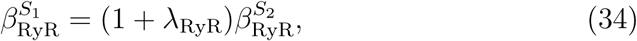

where superscript *S*_1_ and *S*_2_ denote the *S*_1_ and *S*_2_ versions of the parameters, respectively.

##### Passive diffusion fluxes between compartments (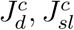**, and** 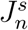)

Following the approach in e.g. [42], diffusion between compartments is considered to occur, on average, between the center of adjacent compartments. Fick’s law of diffusion may then be approximated as

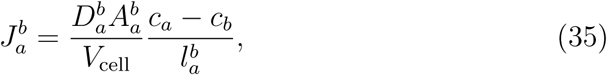

where 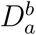 is the diffusion coefficient (in dm^2^/ms) representing the ease with which Ca^2+^ ions flow between the compartments, 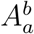 is the area (in dm^2^) of the interface between the compartments, *c*_*a*_ and *c*_*b*_ are the Ca^2+^ concentrations of the compartments (in mM), and 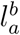 is the distance between the centers of the two compartments (in dm). Again, *V*_cell_ is the total cell volume (in L), and the flux 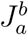 is defined as the number of ions flowing between the compartments per millisecond per total cell volume. Again, in order to reduce the number of parameters, we define the lumped parameter

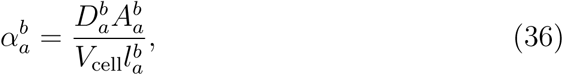

and write the flux as

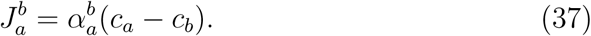

We consider passive diffusive fluxes of this form between the dyad and the bulk cytosol, between the SL and the bulk cytosol, and between the nSR and jSR, and define these fluxes as:

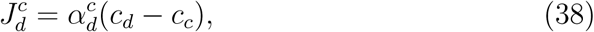

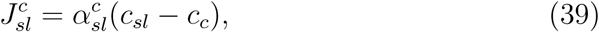

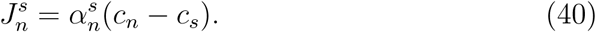

##### Scaling the diffusive fluxes

In the same manner as above, we define adjustment factors 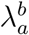 for the diffusive fluxes on the form

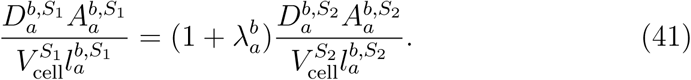

Here, 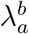 may represent a change in any of the geometrical properties 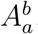, *V*_cell_ or 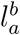, a change in the diffusion coefficient 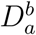, or a combination of these changes. This adjustment is represented in the model for each of the diffusive fluxes by

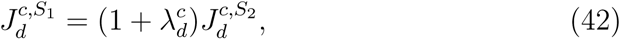

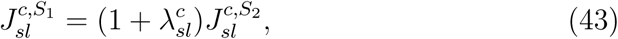

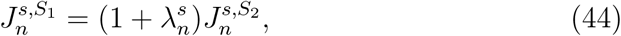

where 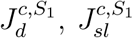, and 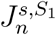 denote the *S*_1_ fluxes and 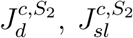, and 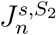 denote the *S*_2_ fluxes.

##### Buffer fluxes (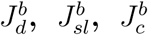, and 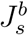)

The chemical reaction between Ca^2+^ and a buffer may be written as

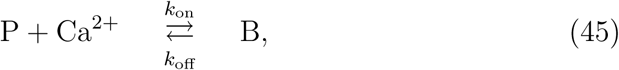

where P represents the buffering protein and B represents Ca^2+^ bound to the buffer. Here, *k*_on_ and *k*_off_ are the rates of the reaction and are given in units of ms^−1^mM^−1^ and ms^−1^, respectively. If we let *B*_tot_ denote the total buffer concentration in some compartment, *c* denote the concentration of free Ca^2+^ and *b* denote the concentration of Ca^2+^ bound to the buffer, the law of mass action (see e.g., [43]) gives that the rate of decrease in the free Ca^2+^ concentration in the compartment and the rate of increase in the concentration of Ca^2+^ bound to a buffer due to Ca^2+^ -buffer reactions is

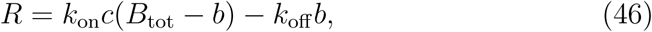

in units of mmol/ms per compartment volume, 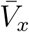 The corresponding flux in terms of mmol/ms per total cell volume, *V*_cell_, may be defined as

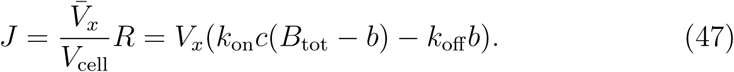

Consequently, the flux of free Ca^2+^ binding to a buffer in the dyad, the SL, the bulk cytosol and the jSR are given by

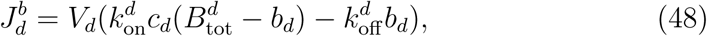

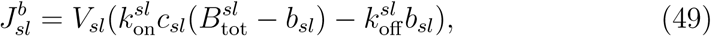

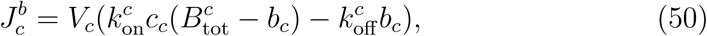

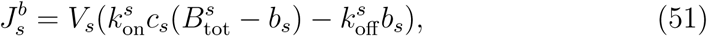

respectively.

##### Scaling the Ca^2+^ buffers

Like for membrane and SR membrane channels, we assume that cells of different levels of maturity contain the same types of Ca^2+^ buffers, with the same rates *k*_on_ and *k*_off_, but that the concentration of the Ca^2+^ buffers, *B*_tot_, may differ for different types of cells. Therefore, we define scaling parameters for the buffer concentrations on the form

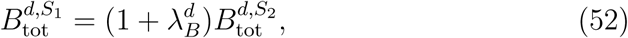

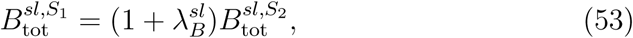

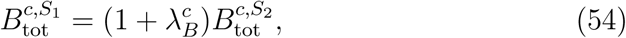

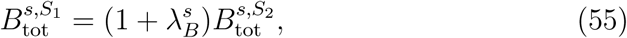

Where 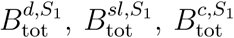, and 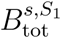 are the buffer concentrations in the *S*_1_ model, and 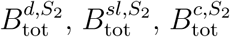, and 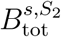 are the buffer concentrations in the *S*_2_ model.

##### Membrane fluxes (*J*_CaL_, *J*_bCa_, *J*_pCa_, and *J*_NaCa_)

The membrane fluxes *J*_CaL_, *J*_bCa_, *J*_pCa_, and *J*_NaCa_ may be defined from the expressions for the corresponding membrane currents, *I*_CaL_, *I*_bCa_, *I*_pCa_, and *I*_NaCa_. Recall from Section 2.2 that the membrane currents are expressed on the form

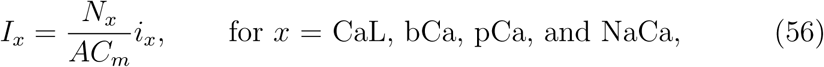

where *N*_*x*_ is the total number of channels of type *x* on the cell membrane, *A* is the total membrane area (in *µ*m^2^), *C*_*m*_ is the specific membrane capacitance (in pF/*µ*m^2^) and *i*_*x*_ is the average single-channel current through a channel of type *x* (in pA). The corresponding membrane fluxes per total cell volume may similarly be defined as

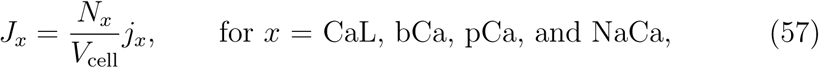

where *V*_cell_ is the total cell volume (in L), and *j*_*x*_ is the average Ca^2+^ flux through a single channel of type *x* (in mmol/ms). The average flux of Ca^2+^ through a single Ca^2+^ channel may be written as

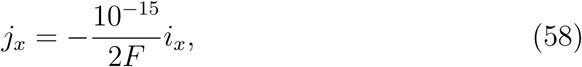

where *F* is Faraday’s constant (in C/mmol), representing the electric charge per mmol of ions with elementary charge. Note that the reason for the factor two in the denominator is that the valence of a Ca^2+^ ion is two, and the factor 10^−15^ is included in the numerator to convert the flux from unit amol/ms to unit mmol/ms. Moreover, the reason for the negative sign in (58) is that the positive direction of the single channel current by convention is from the inside to the outside of the cell, whereas the positive direction of the Ca^2+^ flux is defined to be from the outside to the inside of the cell. Note also that since the Na^+^-Ca^2+^ exchanger exchanges three Na^+^ ions for one Ca^2+^ ion, the flux of one Ca^2+^ ion through the exchanger represents the exchange of one charge instead of two, and a positive current out of the cell is associated with a flux of Ca^2+^ into the cell. Therefore (58) is replaced by

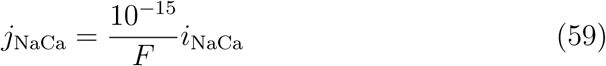

in this case.

Combining (56)–(59), we see that the total membrane fluxes may be written as

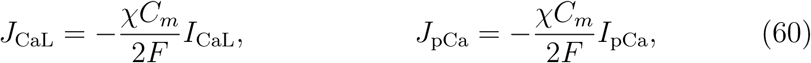

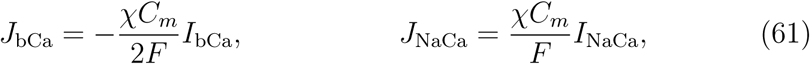

where

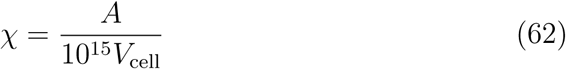

is the surface-to-volume ratio of the cell (in *µ*m^−1^). The expressions for the currents *I*_CaL_, *I*_bCa_, *I*_pCa_, and *I*_NaCa_ are defined in the Supplementary information.

##### Scaling the surface-to-volume ratio

As explained above, we assume that the density of the membrane channels responsible for the Ca^2+^ fluxes may be different for IM and M cells. This change in channel density is represented in the model by scaling the currents (see (4)–(5)), which will also affect the corresponding Ca^2+^ fluxes (60)–(61).

In addition, we again assume that the geometry of the cells (i.e., the membrane area, *A*, and the cell volume, *V*_cell_) may be different for different levels of maturity. From (60) –(63), we see that this change in geometry may be represented by scaling the surface-to-volume ratio, *χ*, by

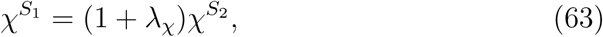

where 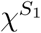 is the surface-to-volume ratio for maturity stage 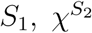 is the value for maturity stage *S*_2_, and *λ*_*χ*_ is an adjustment factor for the surface-to-volume ratio.

### 2.4 Inversion of optical measurements

The inversion procedure, used to construct base model representations of data obtained from optical measurements of the AP and Ca^2+^ transient of hiPSC-CMs, is described below. First, in Section 2.4.1, we describe how optical measurements of hiPSC-CMs are obtained. Next, in Section 2.4.2, we describe how adjustment factors *λ* are set up to represent control (non-drugged) cells from different data sets. In Section 2.4.3, we describe how the effect of a drug is modeled using IC50 values and corresponding factors, denoted by *ε*. The aim of the inversion procedure is to find optimal parameter vectors *λ* and *ε* so that the model parameterized by *λ* and *ε* aligns to the measured data as best possible. This is explained in more detail in Section 2.4.4. In Section 2.4.5, we describe the cost function constructed to measure the difference between the model and the data. Finally, in Section 2.4.6, we describe the continuation-based minimization method used to minimize the cost function in our computations.

#### 2.4.1 Optical measurements

Using previously developed techniques [10], cardiac microphysiological systems were loaded and matured prior to drug exposure. On the day upon which studies were performed, freshly measured drugs (Nifedipine, Lidocaine, Cisapride, Flecainide, and Verapamil) were dissolved into DMSO or media and serially diluted. Each concentration of the drug to be tested was preheated for 15-20 min in a water bath at 37 °C and subsequently sequentially injected in the device. At each dose, after 5 min of exposure, the drug’s response on the microtissue was recorded using a Nikon Eclipse TE300 microscope fitted with a QImaging camera. Fluorescent images were acquired at 200 frames per second using filters to capture GCaMP and BeRST-1 fluorescence as previously described. Images were obtained across the entire chip for 6-8 seconds, with cell excitation paced at 1 Hz, to capture multiple beats for processing.

Fluorescence videos were analyzed using custom Python software utilizing the open source Bio-Formats tool to produce characteristic AP and Ca^2+^ waveforms for each chip and tested drug dose. Briefly, for each analysis, the fluorescent signal was averaged over the chip. The signal was then smoothed using a 3 point median filter, and 5-7 individual action potentials or calcium transients were overlayed by aligning the maximum dF/dt and then averaged into a single transient. For each drug escalation study, we chose the single series with the most continuity between control cases and subsequent drug doses for both AP and Ca^2+^ transient for inversion and mapping analysis.

#### 2.4.2 Definition of adjustment factors

In order to make base model representations of control cells from different data sets, we must define adjustment factors *λ* for a base model parameter set. These adjustment factors represent alterations of the channel densities and geometry of the cells under consideration, as explained above. For example, for each membrane channel type *x*, the adjustment factor *λ*_*x*_ is defined as

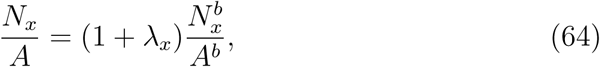

where 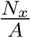 is the channel density on the cell membrane for the fitted model and 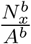 is the channel density in the default base model. We generally consider adjustment factors for the membrane channel densities for all the currents of the model, i.e. *λ*_Na_, *λ*_NaL_, *λ*_CaL_, *λ*_to_, *λ*_Kr_, *λ*_Ks_, *λ*_K1_, *λ*_NaCa_, *λ*_NaK_, *λ*_pCa_, *λ*_bCl_, *λ*_bCa_, and *λ*_f_, although some of the factors are fixed in some cases (see Section 3.1.4).

For the density of an intracellular channel type *x*, the adjustment factor *λ*_*x*_ is similarly defined as

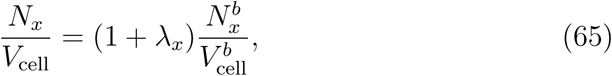

where 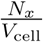 and 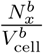 are the channel densities for the fitted model and the default base model, respectively. We consider the following adjustment factors for the intracellular channel densities: 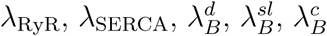, and 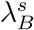. In addition, we consider adjustments to the intracellular diffusion coefficients, 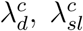, and 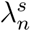 (see (41)). In order to reduce the number of free parameters to be determined in the inversion procedure, we assume that the buffer concentrations change at the same rate in all intracellular compartments, so that we only consider a single adjustment factor

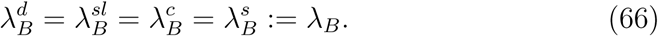

Similarly, we assume that the intracellular diffusion coefficients change at the same rate, so that

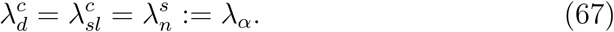

Furthermore, because we wish to avoid ending up with unrealistic values of the surface-to-volume ratio, *χ*, we assume that the scaling factor for the cell surface-to-volume ratio varies little between data sets and only employ two different values of *χ* in the computations. We use the value *χ* = 0.6 *µ*m^−1^ for mature cells and the value *χ* = 0.9 *µ*m^−1^ for immature cells, based on the values used in the Grandi et al. AP model for mature cells [19] and the Paci et al. AP model for hiPSC-CMs [28].

#### 2.4.3 IC50 modeling of drug effects

Following previous modeling of channel blockers (see e.g., [29, 44, 45, 46]), we model the dose-dependent effect of a drug by scaling the channel conductances according to

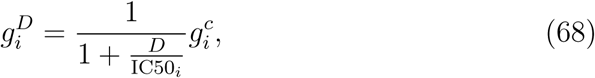

where 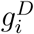 is the conductance of channel *i* in the presence of a drug with concentration *D*, IC50_*i*_ is the drug concentration that leads to 50% block of channel *i*, and 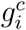 is the channel conductance in the control case (i.e., in the absence of drugs). Specifically, this means that if the drug concentration *D* equals the IC50 value, we have 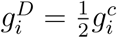.

It should be mentioned that a drug may certainly affect a channel in a more complex manner than is assumed here. The effect of drugs can realistically be represented by introducing new states in Markov models of the ion channel. In such models, the transition rates between different model states are able to represent the properties of drugs (see e.g., [47, 48, 49, 50]). Although Markov model representations of drug effects are more versatile and realistic than the simple blocking assumption employed here (68), it would greatly increase the complexity of the inversion process, as many more parameters would have to be computed.

From (68), we see that for a given drug dose *D >* 0, the effect of the drug would increase if the IC50 value were decreased, and the effect of the drug would be very small if the IC50 value were much larger than the considered dose. In the continuation-based minimization method applied in our computations (see Section 2.4.6 below), it is most practical to deal with parameters that are small when no change occurs and large when large changes occur. Therefore, we introduce the parameters

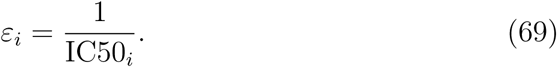

Here, a small value of *ε*_*i*_ represents small effects of a drug while a large value of *ε*_*i*_ represents large effects, and channel blocking is given by

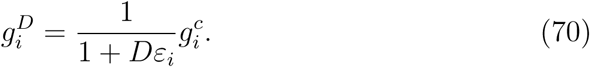

In our computations, we assume that the considered drugs block either *I*_CaL_, *I*_NaL_, *I*_Kr_, or a combination of these currents, and we therefore only consider the *ε*-parameters *ε*_CaL_, *ε*_NaL_, *ε*_Kr_.

#### 2.4.4 Coupled inversion of data from several doses

The control data obtained from different optical experiments tends to vary significantly, and in order to be able to accurately estimate the drug effect from these measurements, the *λ*-parameters must be tuned so that the control model fits the control data as best possible. In addition, we want the *λ* parameters to be constructed such that that the scaling (70) for *ε*_CaL_, *ε*_NaL_, and *ε*_Kr_ is sufficient to fit the model to the drug doses under consideration. In order to increase the chance of obtaining such a control model, we fit the control parameters, *λ*, and the drug parameters, *ε*, simultaneously, instead of first finding the optimal control parameters, *λ*, by fitting the base model to the control data, and then subsequently finding appropriate drug parameters, *ε*, for each dose. In addition, all doses are included in the inversion, so that the estimated values of *ε* are based on *all* the drug doses included in the data set.

In order to illustrate the role of the *λ*- and *ε*-parameters more clearly, consider a simplified model consisting of just two currents, and assume that the base model is given by (see Section 2.2)

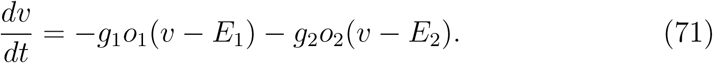

Assume further that we have data from cells with both no drug present and with different doses of a drug (e.g., one low dose and one high dose). We assume that the drug may block any of the two model currents. In the inversion procedure, we try to find optimal values of the four parameters *λ*_1_, *λ*_2_, *ε*_1_ and *ε*_2_ so that the adjusted model of the form

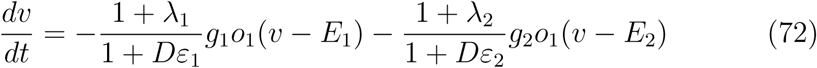

fits the data both for the control case (*D* = 0) and for the considered drug doses. In other words, for a given parameter set *λ*_1_, *λ*_2_, *ε*_1_ and *ε*_2_, we need to compute the solution of the model (72) both for the control case (*D* = 0) and for the considered drug doses and compare the obtained solutions to the corresponding experimental data.

The more general case considered in our computations is conceptually identical; however, as we also consider scaling of parameters that are not assumed to possibly be affected by the drug, we also have some parameters simply scaled by a factor (1+*λ*_*i*_) instead of by 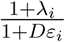.

#### 2.4.5 Properties of the cost function

In order to find the optimal parameters for fitting the model to data, we need to define a cost function that measures the difference between a given model solution and the data. This cost function is defined as

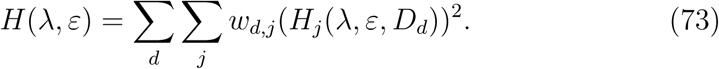

Here, *d* runs over each of the considered drug doses, *D*_*d*_, including the control case (*D*_0_ = 0), and *j* runs over each cost function term, *H*_*j*_, representing various differences between the data and the model solution. The parameters *w*_*d,j*_ represent weights for each of the cost function terms for each of the doses. The weights are generally set to 1, unless otherwise specified.

The considered data from the hiPSC-CM microphysiological systems are traces measuring the membrane potential and the cytosolic Ca^2+^ concentration. As these measurements are optically obtained using voltage- and Ca^2+^ - sensitive dyes, some characteristics of the AP and Ca^2+^ transient cannot be sampled directly from the data, for instance the maximum and minimum values of the membrane potential and Ca^2+^ concentration. However, other biomarkers, like the AP duration, are readily obtained. When necessary for comparing simulation results (with units) and experimental data (unitless), experimental data values are mapped so that the maximum and minimum values of the membrane potential and the Ca^2+^ transient match the maximum and minimum values of the model solution.

Each of the cost function terms, *H*_*j*_, are defined below. The definition of some of the quantities involved in the cost function terms are illustrated in Figure S1 in the Supplementary information.

##### Action potential and Ca^2+^ transient durations

The terms in the cost function include terms for the differences in the AP and Ca^2+^ transient durations of the form

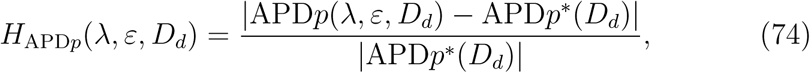

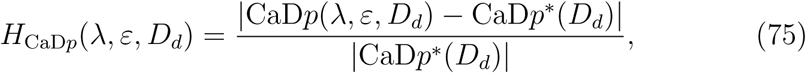

for *p* = 20, 25, …75, 80. Here, as an example, APD30 is defined as the time from the membrane potential is 30% below its maximum value during the upstroke of the AP, *t*_1_, to the time at which the membrane potential again reaches a value 30% below its maximum value during the repolarization phase, *t*_2_. APD30(*λ, ε, D*_*d*_) is the value obtained from the solution of the model given by the parameter vectors *λ* and *ε* for the drug dose *D*_*d*_, while APD30^*^(*D*_*d*_) is the value obtained from data for the drug dose *D*_*d*_. The same notation, with a ‘*’ marking the measured data values, is used for all the terms in the cost function. The Ca^2+^ transient durations, CaD*p*, are defined in the same manner as the AP durations.

##### Integral of the membrane potential

Because the APD*p* values for low values of *p* may be difficult to obtain from the optical measurements due to noise, we also include a term that considers the integral of the membrane potential from *t*_1_ to *t*_2_ (see Figure S1 in the Supplementary information). This term is defined as

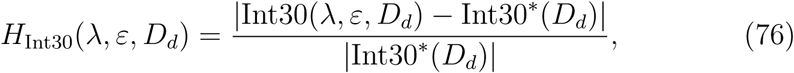

where Int30 is defined as

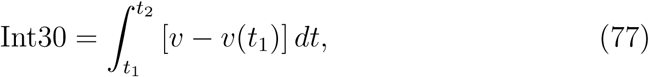

and *v* is the membrane potential. Note that the values of *t*_1_ and *t*_2_ are here the ones defined in the computation of APD30.

##### Norm of the Ca^2+^ transient difference

As there is typically less noise in the data obtained from optical measurements of the Ca^2+^ transient as compared to optical measurements of the membrane potential, we also include a term for the discrete *l*^2^-norm of the difference between the Ca^2+^ transient of the data and the model,

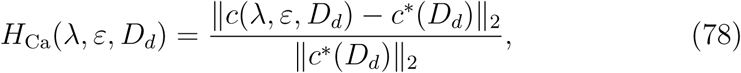

where *c* is the cytosolic Ca^2+^ concentration. When the data *c*^*^ is obtained from optical measurements, the timing of the Ca^2+^ transient relative to the stimulation time is not known. Therefore, the value of *H*_Ca_ is taken as the smallest value obtained when the timing of *c*^*^(*D*_*d*_) may be adjusted to fit the timing of *c*(*λ, ε, D*_*d*_).

##### Upstroke velocity

In order to capture information about the upstroke of the AP and Ca^2+^ transient, we also consider the terms

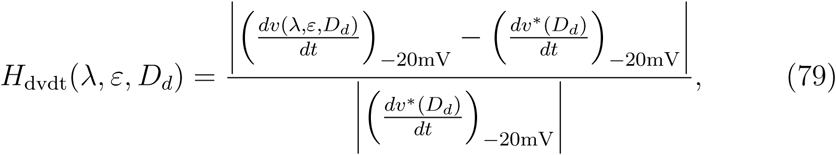

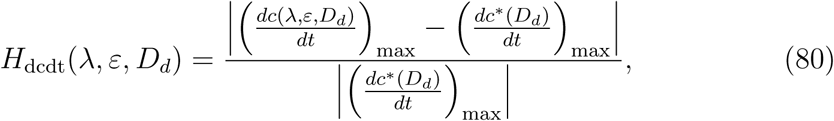

where 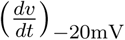 is the upstroke velocity of the AP obtained at *v* = −20 mV, and 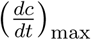 is the maximal upstroke velocity of the Ca^2+^ transient. We use the upstroke velocity obtained at *v* = −20 mV instead of the maximal upstroke velocity to ensure that the value obtained in the model is not determined by the stimulus current. Note, however, that because of the noise in the optical measurements of the membrane potential, the *H*_dvdt_-term is currently only included in the inversions used to determine a mature base model. In that case, the “experimental” data used for parameterization are generated by simulations and therefore *dv/dt* can accurately be computed.

##### Ca^2+^ transient amplitude

Because the Ca^2+^ transient amplitude is one of the main characteristics permitting distinction between block of *I*_CaL_ as opposed to *I*_NaL_, we also include the term

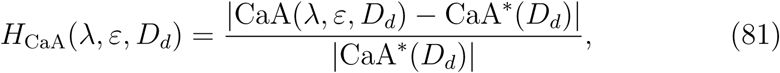

where CaA denotes the Ca^2+^ transient amplitude (see Figure S1 in the Supplementary information).

Note, however, that the actual values of the Ca^2+^ transient amplitude are not known from the optical measurements, and only the relative differences of the amplitude between the control case and the different drug doses are known. Therefore, we do not include the *H*_CaA_-term for the control case. For the non-zero drug doses, we define the data values CaA^*^(*D*_*d*_) so that the relative difference between CaA^*^(*D*_*d*_) and the amplitude in the control model is the same as the relative difference between the amplitude in the data for drug dose *D*_*d*_ and the control data. In other words, CaA^*^(*D*_*d*_) is defined as

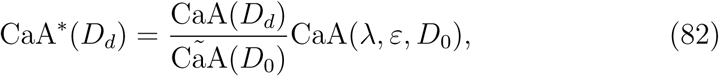

where CãA(*D*_*d*_) and CãA(*D*_0_) are the unitless measured Ca^2+^ transient amplitudes for the drug dose *D*_*d*_ and the control case (*D*_0_ = 0), respectively. Furthermore, CaA(*λ, ε, D*_0_) is the amplitude of the Ca^2+^ transient in the current control model given by the adjustment parameters *λ*.

##### Maximum and resting values of the membrane potential and Ca^2+^ concentration

Where it is desirable to include information about the resting and maximum values of the membrane potential and/or the cytosolic Ca^2+^ concentration, we include terms of the form

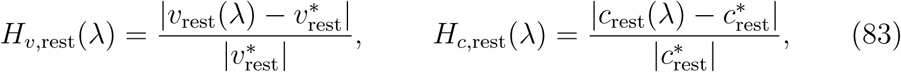

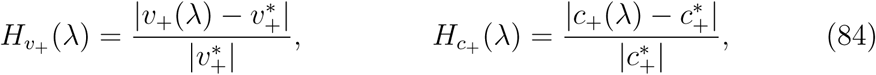

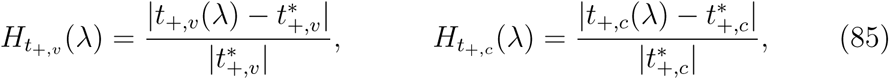

where *v*_rest_ and *c*_rest_ are the resting membrane potential and Ca^2+^ concentration, respectively, defined as the values obtained 10 ms before stimulation in the applied stimulation protocol. Similarly, *v*_+_ and *c*_+_ are the maximum values of the membrane potential and Ca^2+^ concentration, respectively, and *t*_+,*v*_ and *t*_+,*c*_ are the points in time at which these values are reached. Note that these terms are only included when the base model is fitted to the Grandi et al. model to define a mature base model.

##### Information about individual currents

When the inversion procedure is used to define a default base model for mature and immature cells, information about the individual currents and fluxes is also included. These data are obtained from mathematical models of mature [19] and immature [28] cells, and are represented by cost function terms of the form

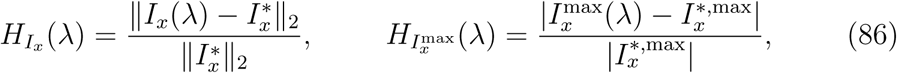

for each of the considered currents or fluxes, *x*. Here, ‖·· ‖_2_ is the discrete *l*^2^-norm, and 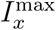 is defined as 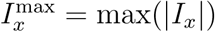.

##### Ca^2+^ balance

We wish to select values of *λ* so that the resulting control model does not exhibit large degrees of drift in the intracellular Ca^2+^ concentrations. Therefore, we include a Ca^2+^ balance term of the form

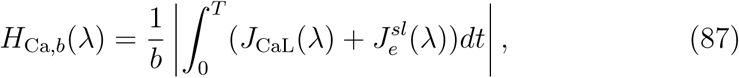

which is zero if the amount of Ca^2+^ entering the cell equals the amount of Ca^2+^ leaving the cell. The main term is here the absolute value of the integral of the sum of the *J*_CaL_ and 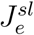 fluxes of the model over the simulated time interval, and *b* is a scaling factor set equal to 0.1 mM in our simulations.

##### Regularization of adjustment factors

In cases where several choices of parameters *λ* and *ε* fit the data equally well, we wish to choose values of *λ* and *ε* close to zero. We therefore include the regularization terms

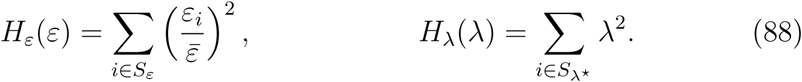

Here, 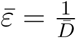, where 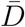 is the median of the non-zero drug doses included in the data set. Furthermore, *S*_*ε*_ is the set of indices for all the individual *ε*-factors, and *S*_*λ*_*\** is the set of indices for the *λ*-values we want to remain as close as feasible to the default base model during the inversion. In the inversions reported below, this set consists of the indices for *λ*_CaL_ and *λ*_Kr_, as the size of these currents is based on measurements of hiPSC-CMs (see below) and because we are especially interested in obtaining reasonable, physiological values for these currents, as we are investigating the drug effects on these currents.

##### Specification of the cost function weights

The choice of terms included in the cost function depends on the specific application of the inversion procedure. In particular, for inversions of data from optical measurements (and for inversions of simulated drugs), we include the terms *H*_APD30_, *H*_APD50_, *H*_APD80_, *H*_CaD20_ − *H*_CaD80_, *H*_int30_, *H*_dcdt_, *H*_Ca_, *H*_*ε*_ and *H*_*λ*_. We only include three APD values, but 13 CaD values as quality of the Ca^2+^ data is generally better than that of the membrane potential data. To make up for the large number of CaD-terms as compared to APD-terms, the weight of the CaD-terms are set to 0.5, while the APD-terms are given the weight 1 (the exception being that the weight of the APD80 and CaD80 terms are each set to 5). The upstroke velocity of the AP is not included because of the high noise level in the membrane potential data.

Furthermore, for the control case, the term *H*_Ca,*b*_ with weight 1 is included, and for the drug doses, *H*_CaA_ is included with weight 10. The large weight in this case is due to the fact that this is one of the most important characteristics for distinguishing between block of *I*_CaL_ and *I*_NaL_. We also include the regularization terms *H*_*ε*_ and *H*_*λ*_ with weights, 0.01 and 10, respectively. All the *ε*-parameters are included in the *ε*-regularization, but only *λ*_CaL_ and *λ*_Kr_ are included in the *λ*-regularization as explained above.

In addition, the weight of all the cost function terms (except *H*_Ca,*b*_) are for the control case multiplied by the number of non-zero doses included in the data set. This is done because a good fit for the control model is essential for being able to use the model to estimate drug effects.

In the inversions aiming to define default values for the immature and mature base models, additional terms, e.g., terms for the individual currents, are also included in the cost function. This is specified in more detail in Sections 3.1.1 and 3.1.2.

#### 2.4.6 A continuation-based minimization method

As outlined in previous sections, we wish to adjust the base model to data by finding *λ*- and *ε*-parameters that minimize a cost function of the form (73), measuring the difference between the input data and the model solution. In order to search for the optimal values of *λ* and *ε*, we apply a continuation-based optimization method (see e.g., [32, 33]). Briefly, continuation is used to simplify the solution of equations or of optimization problems by introducing a *θ*-parameterization such that the solution is known for one value of *θ*. Suppose, for instance, that the parameterization is defined such that the solution is known for *θ* = 0 and the problem we want to solve is defined by *θ* = 1. Then the solution at *θ* = 1 can be found by starting at *θ* = 0 and carefully step towards the solution at *θ* = 1. One advantage with this method is that we can start at a solution that we know is correct (at *θ* = 0) and then take *small steps* towards the goal at *θ* = 1. For the problem of inverting membrane potential and Ca^2+^ traces, this method has proven to be useful.

##### Cost function in the continuation case

More specifically, we assume that, for each drug dose, *D*_*d*_, (including the control case) the data we are trying to invert is given by some vector pair (*v*^1^(*D*_*d*_), *c*^1^(*D*_*d*_)), where *v*^1^(*D*_*d*_) is the membrane potential and *c*^1^(*D*_*d*_) is the Ca^2+^ concentration. In addition, from the default base model specified by *λ* = *ε* = 0, we can compute a vector pair (*v*^0^, *c*^0^) for the membrane potential and Ca^2+^ concentration as the starting point of the inversion.

The goal of the continuation method is to compute a path for *λ* and *ε* from *λ* = *ε* = 0, which fit (*v*^0^, *c*^0^) perfectly, to some *λ* and *ε* that fit the final data (*v*^1^(*D*_*d*_), *c*^1^(*D*_*d*_)) for each of the drug doses, *D*_*d*_, as best as possible. This is done by defining a cost function of the form

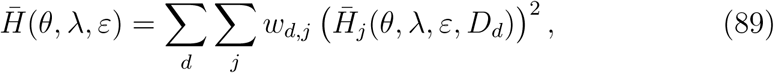

for the intermediate steps in the algorithm. Here, *θ* is a parameter that is gradually increased from zero to one. In the definition (89), the terms 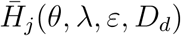 correspond to each of the terms *H*_*j*_(*λ, ε, D*_*d*_) defined above.

Specifically, the terms take the form

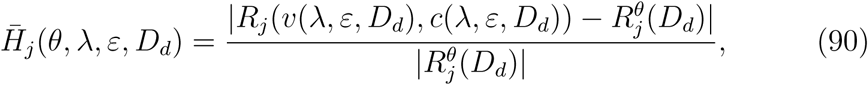

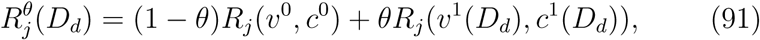

where *R*_*j*_(*v, c*) represent different characteristics of the AP or Ca^2+^ transient, e.g., the AP duration at some percentage or the upstroke velocity (see (75)– (86) ^1^). In the case 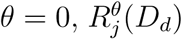 is equal to the terms defined by the default model (*λ* = *ε* = 0) for all the doses *D*_*d*_. Therefore, 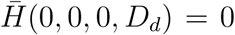,^2^ so the optimal solution for *θ* = 0 is *λ* = *ε* = 0. In the case *θ* = 1, the terms 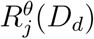 are equal to the characteristics computed for the data we wish to invert. In other words, 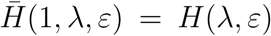, where *H*(*λ, ε*) is defined in (73). For the intermediate values of *θ*, the characteristics 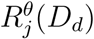 represent weighted averages of characteristics of the model used as a staring point for the inversion (*λ* = *ε* = 0) and the data we are trying to invert. Therefore, we expect the optimal values of *λ* and *ε* to gradually move from zero to the optimal values for the data as *θ* is increased from zero to one.

##### The minimization algorithm

In the minimization algorithm, we find the optimal solution in *M* iterations. We define *θ*_*m*_ = Δ*θ*·*m* for *m* = 0, …, *M* where Δ*θ* = 1*/M*. For *m* = 1, …, *M*, we assume that the optimal values *λ*(*θ*_*m*−1_) and *ε*(*θ*_*m*−1_) have been computed, and we want to find *λ*(*θ*_*m*_) and *ε*(*θ*_*m*_) by finding the minimum of 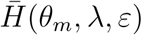. Since the step in *θ* is small, we assume that the changes in *λ* and *ε* are also relatively small. We use the Nelder-Mead algorithm [51] to minimize 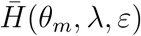, and we use *λ*(*θ*_*m*−1_) and *ε*(*θ*_*m*−1_) as suggestions for the starting vectors to find *λ*(*θ*_*m*_) and *ε*(*θ*_*m*_). However, in order to increase the chance of finding the true optimal value in every iteration, we start the Nelder-Mead algorithm from several randomly chosen starting vectors in the vicinity of *λ*(*θ*_*m*−1_) and *ε*(*θ*_*m*−1_). Figure S2 in the Supplementary information illustrates the development of the *ε*-values in an inversion aiming to characterize a drug.

##### Technical specifications

In the applications presented below, we use *M* = 20, and in each iteration *m*, we draw 63 guesses (as the specific computer used for these simulations has 64 cores) for the starting vectors for the Nelder-Mead algorithm from [*λ*(*θ*_*m*−1_)−0.2, *λ*(*θ*_*m*−1_)+0.2] and 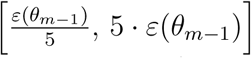 for *λ* and *ε*, respectively. In the first 15 iterations, we use five iterations of the Nelder-Mead algorithm for each guess, and for the last five iterations we use 25 iterations of the Nelder-Mead algorithm. For each new parameter set, we generally run the simulation for 15 AP cycles before measuring the AP and Ca^2+^ transient, unless otherwise specified.

### 2.5 Identifiability of the base model based on singular value decomposition of currents

In the inversion procedure outlined above, we try to find the optimal adjustment factors *λ* and *ε* for the model so that the AP and the cytosolic Ca^2+^ transient in the model solution match measurements of the AP and Ca^2+^ transient as best possible. An important element to consider in this process is whether the identified adjustment factors found by the inversion procedure are the only combination of adjustment factors that fit the data, or whether other adjustment factors might exist which fit the data equally well.

In order to investigate the identifiability of the adjustment factors for the currents in the base model, we apply a method based on a singular value decomposition (see e.g., [52, 53]) of the currents. This approach is described in detail in [54]. In short, the identifiability of the currents is investigated by collecting the model currents at time points *t*_*n*_ = *n*Δ*t*, for *n* = 1, …, *N*_*t*_ into a matrix 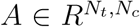, where *N*_*c*_ is the number of model currents. Then, the singular value decomposition of the matrix

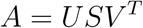

is computed. Here, the matrices 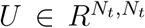 and 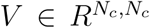 are unitary matrices, and the matrix 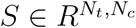 is a diagonal matrix with singular values *σ* _*i*_ along the diagonal. The columns *u*_*i*_ and *v*_*i*_ of *U* and *V*, respectively, are the associated singular vectors.

From the properties of the singular value decomposition it can be shown that perturbations of the adjustment factors along singular vectors *v*_*i*_ associated with large singular values *σ* _*i*_ are expected to result in significant changes in the AP, whereas perturbations of the adjustment factors along singular vectors *v*_*i*_ associated with small singular values are, accordingly, expected to result in small changes in the AP.

In [54] it was shown that this expected result seemed to hold in the case of three well-known AP models of mature ventricular cardiomyocytes. In addition, it was demonstrated how this analysis could be used to define an *identifiability index* for individual model currents. This index was defined for each current *j* = 1, …, *N*_*c*_ as

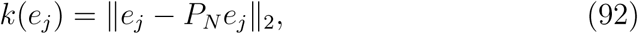

where 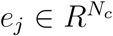 is the vector that is one in element number *j* and zero elsewhere. Moreover, 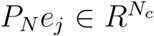 is the projection of *e*_*i*_ onto the unidentifiable space spanned by the singular vectors *v*_*i*_ associated with small singular values (or small perturbation effects). In other words, if *k*(*e*_*j*_) is close to zero, almost the entire current *I*_*j*_ is in the unidentifiable space, and we cannot be sure that the value of the associated adjustment factors *λ*_*j*_ or *ε*_*j*_ are the only values that fit the data (i.e., result in the same AP). On the other hand, if *k*(*e*_*j*_) is close to one, we expect that other values of *λ*_*j*_ or *ε*_*j*_ would not fit the data as well as the currently assumed values, as perturbations of the adjustment factors would result in large changes in the AP.

Note that this approach only aims to determine the identifiability of the adjustment factors for the membrane currents. The analysis could be extended to include other state variables than just the membrane potential (e.g., the Ca^2+^ concentrations). In this case, the identifiability of the remaining adjustment factors might also be suggested. However, at this stage primary focus is on identifying drug effects on membrane ion channels, so we are principally interested in ensuring that the adjustment factors for the currents are unique.

## 3 Results

Below, we demonstrate a few applications of the method outlined above. First, in Section 3.1, we define the default immature and mature parameterizations of the general base model formulation. We also demonstrate that these models exhibit high gain and graded release of the Ca^2+^ fluxes. In addition, we illustrate the identifiability of the model currents using SVD analysis, as described above. This analysis is used to determine which model currents should be fixed in the applications of the inversion procedure. Next, in Section 3.2, we use the inversion procedure to identify drug effects for data generated by simulations. Finally, in Section 3.3, we apply the inversion procedure to identify drug effects from data obtained from optical measurements of hiPSC-CMs.

### 3.1 The base model

Here, we set up the default mature and immature base model formulations used in the inversion procedure in the following sections.

#### 3.1.1 Base model approximation of the Grandi model

The base model is fitted to approximate the Grandi et al. model using the inversion procedure described above; the right panel of Figure 3 shows the AP and Ca^2+^ transient of the Grandi et al. model [19] for healthy mature ventricular cardiomyocytes and the AP and Ca^2+^ transient of the mature version of the base model. In the inversion, the cost function includes all the terms defined in Section 2.4.5, except for the regularization terms (88). The currents *I*_Na_, *I*_CaL_, *I*_to_, *I*_Kr_, *I*_Ks_, *I*_K1_, *I*_NaCa_, *I*_pCa_, and *I*_bCa_, as well as the fluxes *J*_RyR_ and *J*_SERCA_ are included in (86). All terms measuring the difference in membrane potential or Ca^2+^ concentration are given the weight *w*_*j*_ = 1 and the terms measuring differences in the currents are given the weight *w*_*j*_ = 0.5.

**Figure 3:**
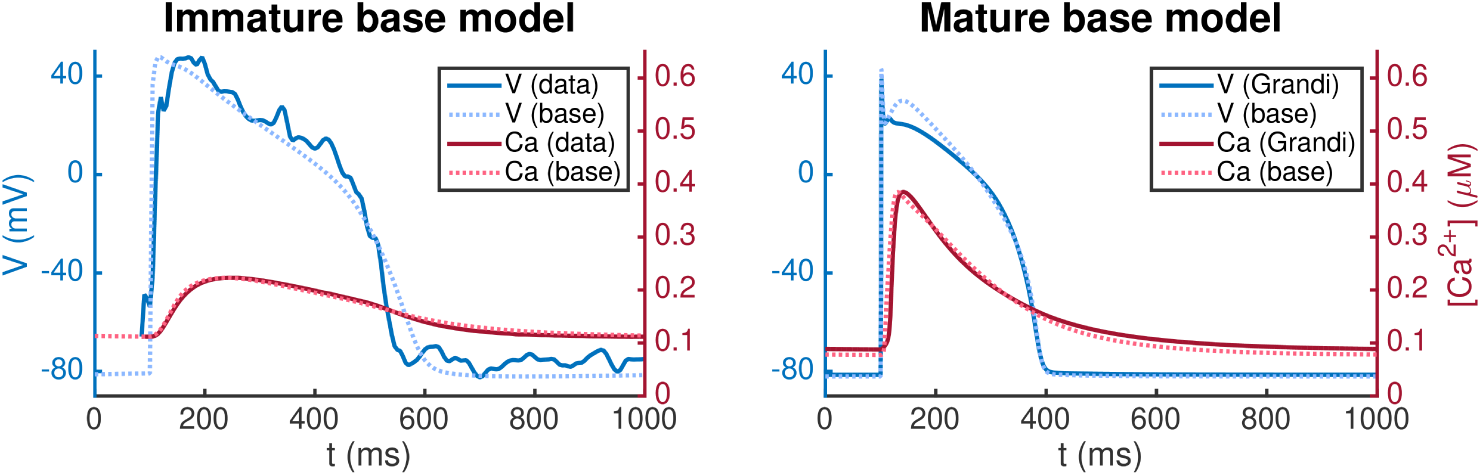
AP (blue) and cytosolic Ca^2+^ transient (red) for the immature and mature versions of the base model. In the left panel, the base model is adjusted to fit data obtained from optical measurements of the AP and Ca^2+^ transient of hiPSC-CMs. In the right panel, the base model is adjusted to approximate the Grandi et al. model [19] of mature cardiomyocytes.

As mentioned above, we define the *default* base model as the mature base model because this model will remain fixed, whereas the immature models will change depending on experimental data. The parameter values obtained in the inversion procedure therefore define the default base model and are specified in the Supplementary information.

#### 3.1.2 Immature base model

The left panel of Figure 3 shows the solution of the immature base model fitted to optical measurements of the AP and Ca^2+^ transient of immature cells. In this case, the cost function consists of the terms 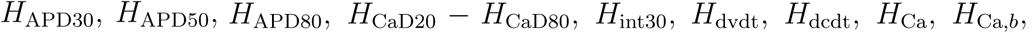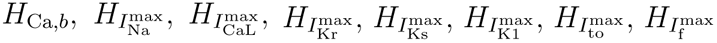, where the information about the currents is obtained from the Paci et al. model [28] which is based on patch-clamp recordings of the ionic currents of hiPSC-CMs from [55]. The terms *H*_CaD20_ − *H*_CaD75_ are given the weight 0.5, and *H*_APD80_ and *H*_CaD80_ are given the weight 5. Furthermore, 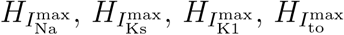, and 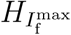 are given the weight 0.5 and 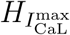 and 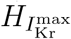 are given the weight 5. The remaining terms are given the weight 1.

The mapping between immature and mature cells returned by the inversion procedure are reported in Table 2. Note that these factors represent the default immature base model to be used as a starting point for the inversion of the remaining control data sets. In other words, the specific adjustment factors between the immature and mature models will differ for each new data set.

**Table 2:** Values defining the maturation map between the default immature and mature base models illustrated in Figure 3. The mature parameters, *p*^*M*^, are related to the immature parameters, *p*^*IM*^, by the relation *p*^*M*^ = (1 + *λ*)*p*^*IM*^. See Sections 2.2–2.3 for more detailed definitions of each of the *λ*-values.

|  |  |  |  |
| --- | --- | --- | --- |
| $\lambda_{\text{Na}}$ | 2.00 | $\lambda_{\text{CaL}}$ | -0.53 |
| $\lambda_{\text{NaL}}$ | -0.08 | $\lambda_{\text{bCa}}$ | 3.90 |
| $\lambda_{\text{Kr}}$ | -0.54 | $\lambda_{\text{pCa}}$ | -0.85 |
| $\lambda_{\text{Ks}}$ | 0.68 | $\lambda_{\text{NaCa}}$ | -0.69 |
| $\lambda_{\text{K1}}$ | 2.23 | $\lambda_{\text{RyR}}$ | -0.12 |
| $\lambda_{\text{to}}$ | 8.45 | $\lambda_{\text{SERCA}}$ | -0.53 |
| $\lambda_{\text{f}}$ | -0.99 | $\lambda_{\text{sl}}^c, \lambda_{\text{n}}^s, \lambda_{\text{d}}^c$ | -0.02 |
| $\lambda_{\text{bCl}}$ | 42.4 | $\lambda_{\text{B}}^c, \lambda_{\text{B}}^d, \lambda_{\text{B}}^{\text{sl}}, \lambda_{\text{B}}^s$ | -0.55 |
| $\lambda_{\text{NaK}}$ | -0.16 | $\lambda_{\chi}$ | -0.33 |

#### 3.1.3 High gain and graded release of the base model

As mentioned above, the base model formulation of Ca^2+^ release is designed to exhibit both high gain and graded release. This has proved impossible to achieve using common pool models (see e.g., [56, 57]), as discussed in more detail in Section 4.2 below. The Ca^2+^ release model we have designed (see (8)–(12) and (26)–(31) above) differs from the classical common pool models in two ways: first, release of Ca^2+^ from the SR is not directed into the dyad (d), but rather directly to the subsarcolemmal (SL) space (see Figure 2), and, second; the release mechanism is formulated in terms of both an availability rate and open probability for the RyRs (see (29)).

In Figure 4, we show that this model exhibits high gain and graded release both when the IM and M parameters are applied. In the figure, we report the peak of the *J*_CaL_ and *J*_RyR_ fluxes as well as the integrated fluxes for simulations in which the membrane potential is fixed at specific values. The remaining state variables of the model start at the default initial conditions corresponding to the default resting membrane potential of the model, and the simulations record the *J*_CaL_ and *J*_RyR_ fluxes resulting from the clamped membrane potential.

**Figure 4:**
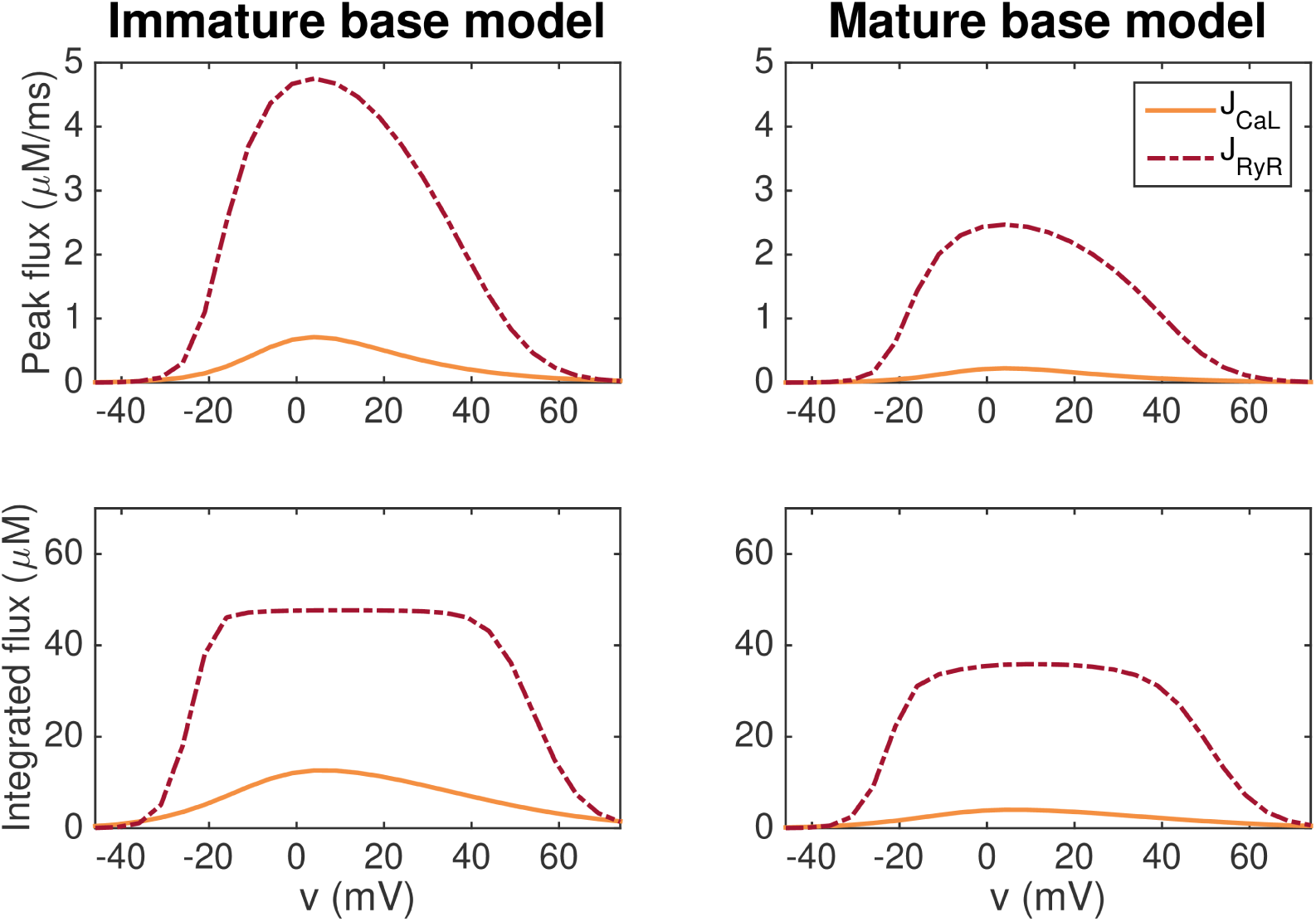
Graded release for the immature (left) and mature (right) versions of the base model. In the upper panel, we report the peak of the *J*_CaL_ and *J*_RyR_ fluxes for simulations in which the membrane potential is fixed at specific values between −50 mV and 80 mV. In the lower panel, we show the fluxes integrated with respect to time from *t* = 0 ms to *t* = 100 ms. After 100 ms, both *J*_CaL_ and *J*_RyR_ have roughly returned to their resting levels.

We observe that for most values of *v*, the *J*_RyR_ flux is considerably larger than the *J*_CaL_ flux, indicating high gain. Furthermore, a small *J*_CaL_ flux seems to be associated with a small *J*_RyR_ flux, whereas a large *J*_CaL_ flux is associated with a large *J*_RyR_ flux, indicating graded release.

#### 3.1.4 Identifiability of the currents in the immature base model

In order to investigate the identifiability of the individual model currents, we apply the singular value decomposition analysis from [54] described in Section 2.5.

In Figure 5, titles above each plot indicate the value of each of the singular values of the current matrix, *A*. The upper plots below the singular values show the singular vectors corresponding to each of the singular values. Here, each letter corresponds to a single current specified in the table on the right-hand side. The below left plots show the values of the cost function (73) evaluated using the default immature base model as data and a perturbed model as the model solution. In the perturbed model, the maximum conductances are perturbed with *λ*-values (see (5)) equal to *ω* · *v*_*i*_, where *v*_*i*_ is the considered singular vector and *ω* is varied between zero and one. The cost function includes the terms *H*_APD30_, *H*_APD50_, *H*_APD80_, and *H*_Int30_ with weight 1 for all terms except *H*_APD80_, which is given the weight 5. The maximum values of *H* are given in the top of the plots. The right plots show the solutions resulting from the perturbations for a few selections of *ω*.

**Figure 5:**
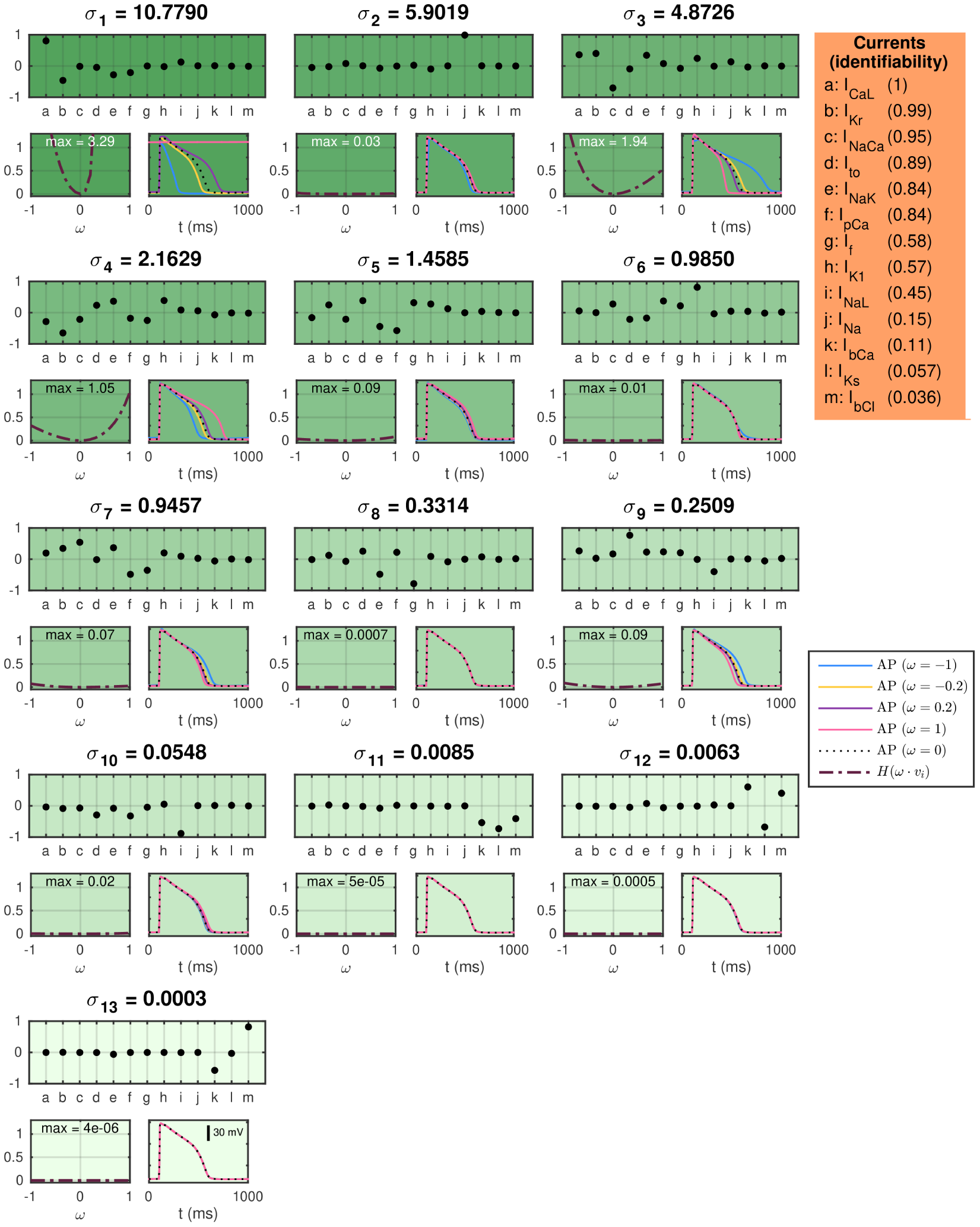
SVD analysis of the currents in the immature base model. The titles above each plot give the singular values of the current matrix *A*, and the upper plots show the corresponding singular vectors. The below plots show how a perturbation of the currents corresponding to the singular vector affects the computed AP for a few examples (right) and measured by a cost function (left). The identifiability index (92) of each current is given in the orange panel.

In [54] it was shown that perturbations of the maximum conductances along singular vectors corresponding to large singular values generally resulted in large perturbation effects, whereas perturbations along singular vectors corresponding to small singular values generally resulted in small perturbation effects for the ten Tusscher et al. [58], the Grandi et al. [19] and the O’Hara et al. [18] AP models. Figure 5 shows correspondence to this result for the immature base model; the main discrepancy is observed for *σ* _2_, which corresponds to a singular vector consisting almost exclusively of the fast sodium current, *I*_Na_. The perturbation effects may be very small for this singular value because the upstroke velocity, physiologically governed by *I*_Na_, is not included in the cost function (cf. [54]).

In order to quantify the identifiability of the individual currents, we compute the identifiability index, *k*, defined in (92). The unidentifiable space is defined as the space spanned by the singular vectors *v*_*i*_ whose maximum value of *H*(*ω*·*v*_*i*_) for 0 ≤ *ω* ≤ 1 is smaller than 0.05. The computed values of the identifiability index for each of the model currents are given in the orange box on the right-hand side of Figure 5. A value of *k* close to 1 indicates a high degree of identifiability, while a value of *k* close to 0 indicates an unidentifiable current.

From the indices in Figure 5, we see that *I*_CaL_, *I*_Kr_, and *I*_NaCa_ are highly identifiable in the immature base model, but that the currents *I*_NaL_, *I*_Na_, *I*_bCa_, *I*_Ks_, and *I*_bCl_ has an identifiability index below 0.5. As a consequence, we fix the conductance of *I*_Na_, *I*_bCa_, *I*_Ks_, and *I*_bCl_ in the applications of the inversion procedure presented below. In addition, we are aware that the *I*_NaL_ current might be hard to identify, and that estimated drug effects for this current are associated with a level of uncertainty (see also [59]).

### 3.2 Identification of drug effects on hiPSC-CMs based on simulated data

Our first application of the inversion procedure is to identify drug effects as based on simulated data. To generate these data, we set *λ*_CaL_ = *λ*_NaL_ = *λ*_Kr_ = 0.1 in the immature version of the base model. In addition, we apply a set of *ε*-values to represent five specific drugs – Nifedipine, Lidocaine, Cisapride, Flecainide, and Verapamil. We assume that Nifedipine is a pure *I*_CaL_-blocker with an IC50 value of 10 nM, that Lidocaine is a pure *I*_NaL_-blocker with an IC50 value of 10 *µ*M, and that Cisapride is a pure *I*_Kr_-blocker with an IC50 value of 10 nM. Furthermore, Flecainide is assumed to block a combination of all three currents with IC50 values of 25 *µ*M, 20 *µ*M and 10 *µ*M for *I*_CaL_, *I*_NaL_, and *I*_Kr_, respectively. Verapamil is assumed to block *I*_CaL_ with an IC50 value of 200 nM and *I*_Kr_ with an IC50 value of 500 nM. Both when the data is generated and in the inversion procedure, we record the sixth generated AP following each parameter change.

Figure 6 shows the result of the inversion procedure using the *λ*-values *λ*_CaL_, *λ*_NaL_, and *λ*_Kr_ and the *ε*-values *ε*_CaL_, *ε*_NaL_, and *ε*_Kr_ as free parameters in the inversion procedure. The left panel shows the *ε*-values used to generate the data (yellow) and the corresponding *ε*-values returned by the inversion procedure (pink). The center and right panels shows the AP and Ca^2+^ transient, respectively, for the control case and for each of the drug doses included in the data sets. The solid lines show the simulated data and the dotted lines show the solutions generated by the model using the *λ*- and *ε*-values returned by the inversion procedure. Note that to clearly see differences in the Ca^2+^ transient amplitude, the Ca^2+^ transients are adjusted so that the Ca^2+^ transient amplitude is preserved, but the minimum Ca^2+^ concentration is set to zero. We observe that the inversion procedure is able to identify the correct *ε*-values accurately, excepting the *ε*-value for Lidocaine, which is predicted to be considerably lower than the value used to generate the data.

**Figure 6:**
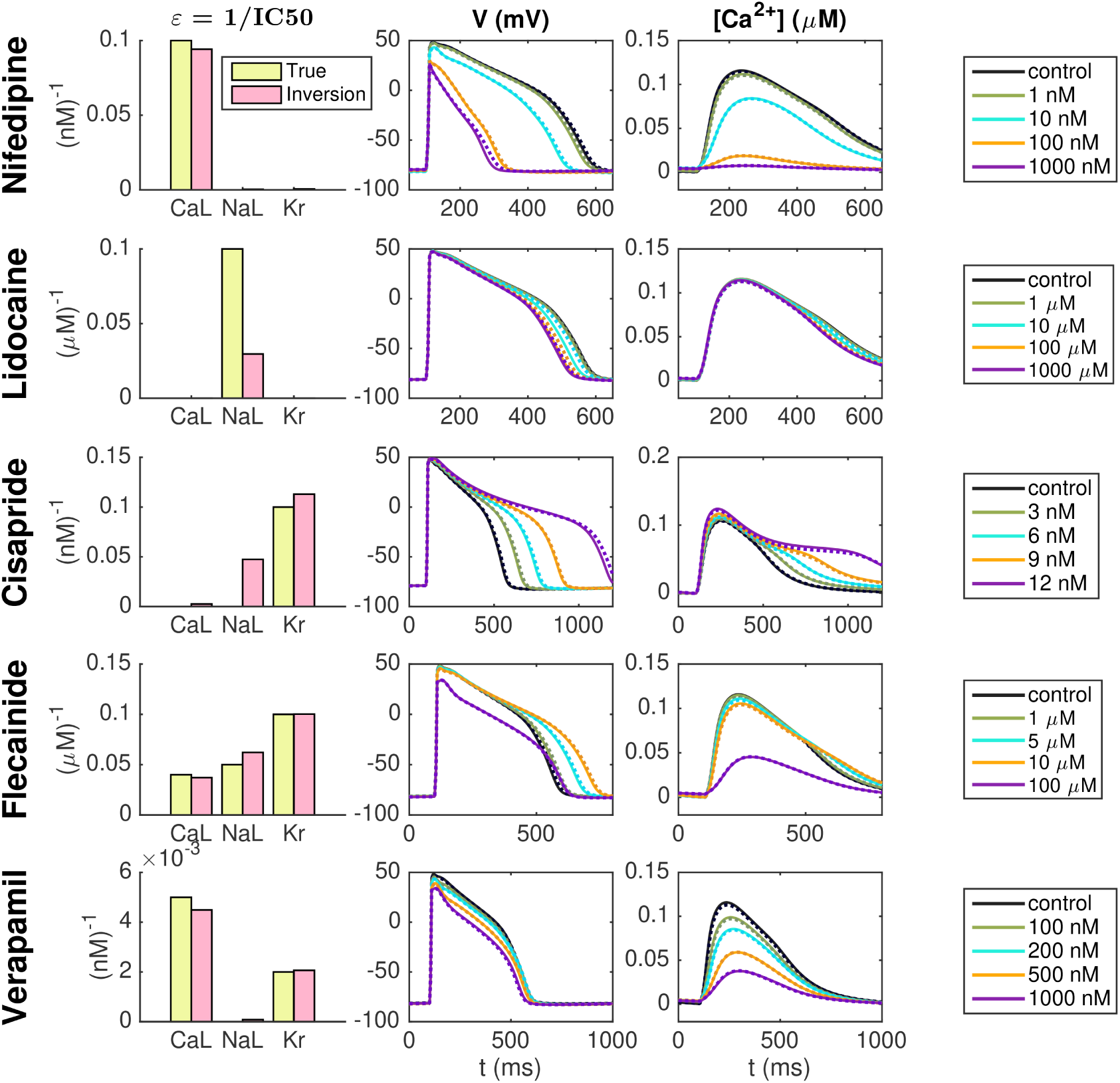
Identification of drug effects for five drugs based on simulated data. The *λ*-values *λ*_CaL_, *λ*_NaL_, and *λ*_Kr_ and the *ε*-values *ε*_CaL_, *ε*_NaL_, and *ε*_Kr_ are allowed to vary in the inversion. The left panel shows the *ε*-values used to generate the simulated drug data (yellow) and the corresponding *ε*-values estimated by the inversion procedure (pink). The center and right panels show the AP and Ca^2+^ transients, respectively, for each of the drug doses included in the data sets. Solid lines represent the simulated data and dotted lines show the fitted model solutions returned by the inversion procedure. Note that to clearly see changes in the Ca^2+^ transient amplitude, the Ca^2+^ transients are adjusted so that the Ca^2+^ transient amplitude is preserved, but the minimum value is set to zero in all cases.

### 3.3 Identification of drug effects on hiPSC-CMs based on optical measurements

Below, we present use of the inversion procedure to identify drug effects on hiPSC-CMs from optical measurements of the AP and Ca^2+^ transient.

#### 3.3.1 Nifedipine

Figure 7 shows the result of the inversion procedure applied to data from optical measurements of hiPSC-CMs exposed to the drug Nifedipine. The data includes the control case with no drug present and four different drug doses (3 nM, 30 nM, 300 nM, and 3000 nM). The left panel of Figure 7A shows the membrane potential and Ca^2+^ traces obtained from optical measurements, and the center panel shows the corresponding solutions of the immature version of the base model fitted to the optical measurements. Note that the values of the data are mapped so that the maximum and minimum values of the membrane potential and Ca^2+^ concentration match those of the fitted immature model. Panel C of Figure 7 compares the experimentally measured data and the fitted model for each of the doses. We observe that the model seems to fit the data quite well for most of the doses, but that the Ca^2+^ transient appears to last a bit longer in the model than in the data for the highest considered drug doses.

**Figure 7:**
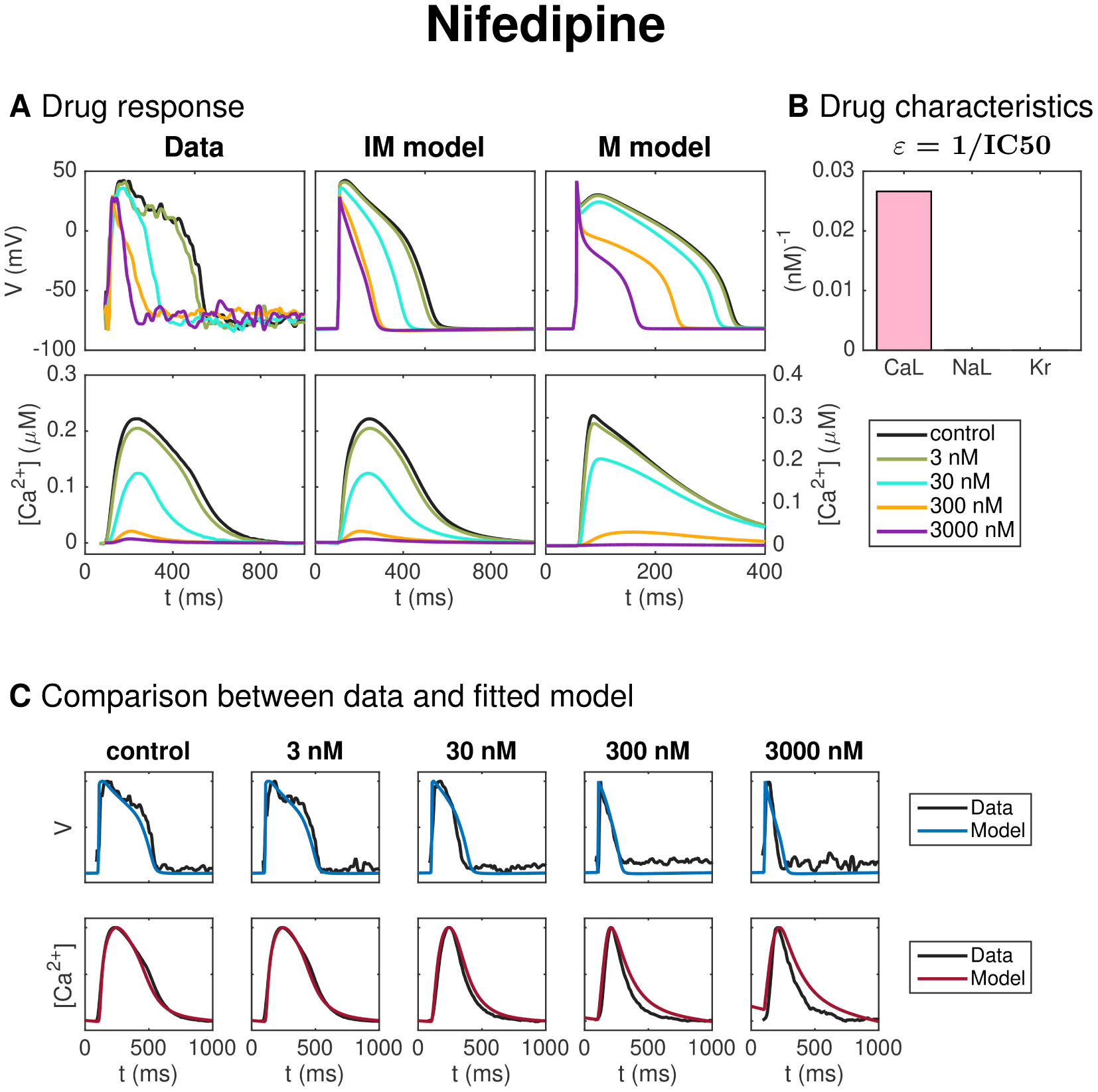
Identification and mapping of drug effects for the drug Nifedipine based on optical measurements of the AP and Ca^2+^ transient of hiPSC-CMs. (**A**) AP and Ca^2+^ transient in the control case and for four drug doses for the data (left) and the fitted immature (IM) model (center). The predicted drug effects for mature (M) cells are given in the right panel (note that the scaling of the axes is adjusted for the M case). Note also that, to clearly see differences in the Ca^2+^ transient amplitude, the displayed Ca^2+^ concentrations are adjusted so that the Ca^2+^ transient amplitude is preserved, but the resting concentration is set to zero in each case. (**B**) Drug effect on *I*_CaL_, *I*_NaL_ and *I*_Kr_ in the form of *ε*-values estimated by the inversion procedure. (**C**) Comparison between the measured membrane potential and Ca^2+^ traces and the fitted immature model solutions for each of the doses in the data set.

The dose-dependent effect of the drug on the *I*_CaL_, *I*_NaL_ and *I*_Kr_ currents are modeled using IC50 values (Section 2.4.3). The values of 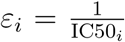 for *i* = CaL, NaL, and Kr are given in Figure 7B. A large value of *ε*_*i*_ corresponds to a large drug effect on the current *i*, and a small value of *ε*_*i*_ corresponds to a small drug effect on the current *i*. From Figure 7B, we observe that the inversion procedure predicts that Nifedipine primarily blocks *I*_CaL_.

The IC50 values corresponding to the estimated *ε*-values for *I*_CaL_, *I*_NaL_ and *I*_Kr_ are given and compared to literature values in Table 3 for all the five considered drugs of this section (see Discussion for a discussion of these results).

**Table 3:** Comparison between the IC50 values obtained from the inversion procedure and values found in literature.

|  |  | Nifedipine | Lidocaine | Cisapride | Flecainide | Verapamil |
| --- | --- | --- | --- | --- | --- | --- |
| CaL | **Inversion**<br>Literature | **38 nM**<br>12 nM [60]<br>60 nM [61] | **3400 μM** | **775 nM**<br>11 800 nM [60] | **9 μM**<br>26 μM [38]<br>27 μM [60] | **495 nM**<br>202 nM [38]<br>200 nM [60]<br>100 nM [62] |
| NaL | **Inversion**<br>Literature | **23 600 nM** | **4.3 μM**<br>11 μM [38] | **120 nM** | **47 μM**<br>19 μM [38] | **23 000 nM** |
| Kr | **Inversion**<br>Literature | **40 200 nM**<br>440 000 nM [60]<br>275 000 nM [63] | **50 000 μM** | **13 nM**<br>12 nM [38]<br>20 nM [60]<br>6.5 nM [64] | **1.9 μM**<br>0.7 μM [38]<br>1.5 μM [60] | **2150 nM**<br>499 nM [38]<br>250 nM [60]<br>143 nM [65] |

#### 3.3.2 Lidocaine

Figure 8 shows similar results for inversion of measurements of hiPSC-CMs exposed to the drug Lidocaine. In panel A, we observe that the AP duration is reduced by the drug, and in panel B, we observe that the inversion procedure predicts that the drug primarily blocks the *I*_NaL_ current.

**Figure 8:**
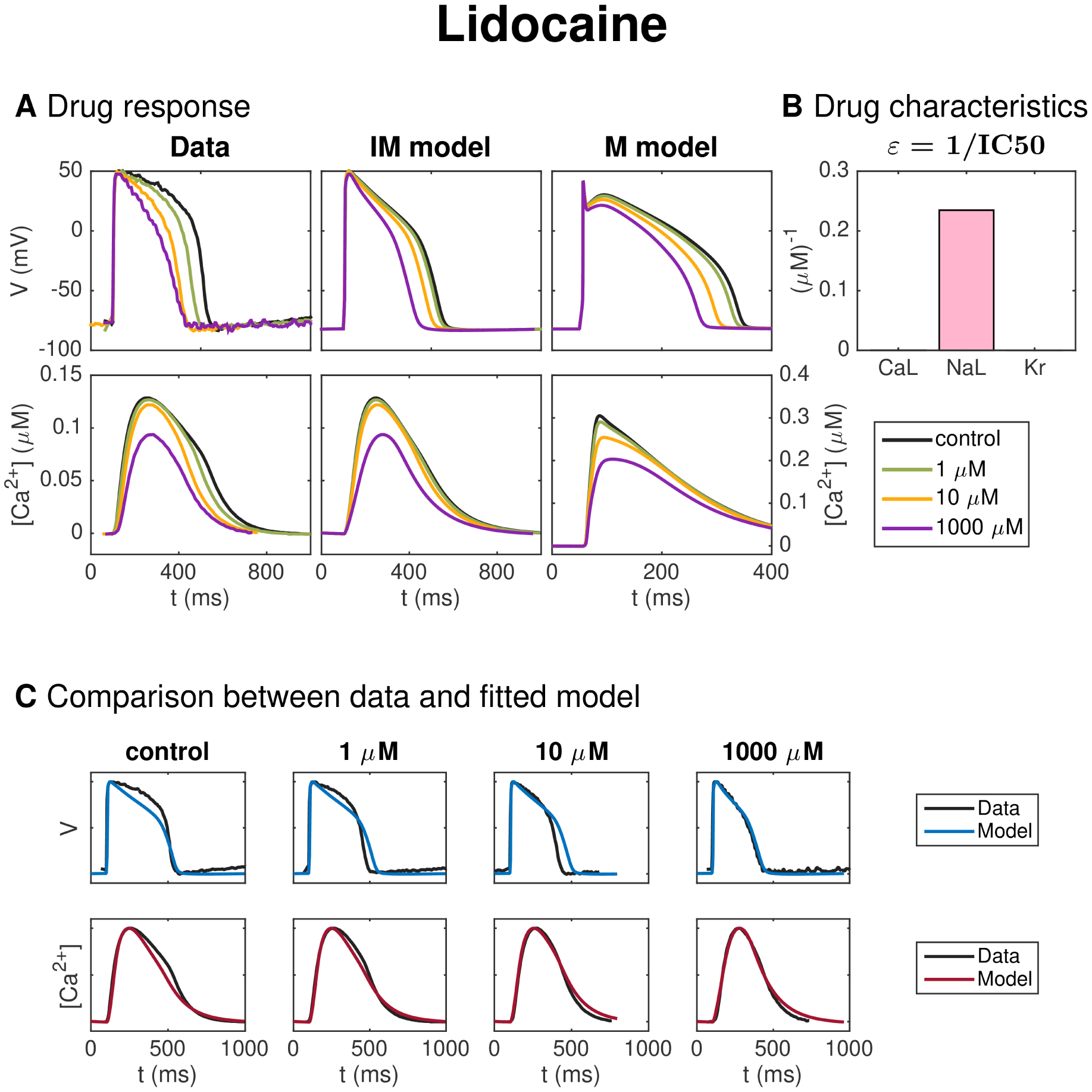
Identification and mapping of drug effects for the drug Lidocaine based on optical measurements of the AP and Ca^2+^ transient of hiPSC-CMs following the same structure as Figure 7.

#### 3.3.3 Cisapride

Figure 9 shows the result of the inversion procedure applied to a data set for hiPSC-CMs exposed to the drug Cisapride. In panel A, we observe that the drug increases the AP duration. In panel B, we observe that the inversion procedure predicts that Cisapride primarily blocks the *I*_Kr_ current.

**Figure 9:**
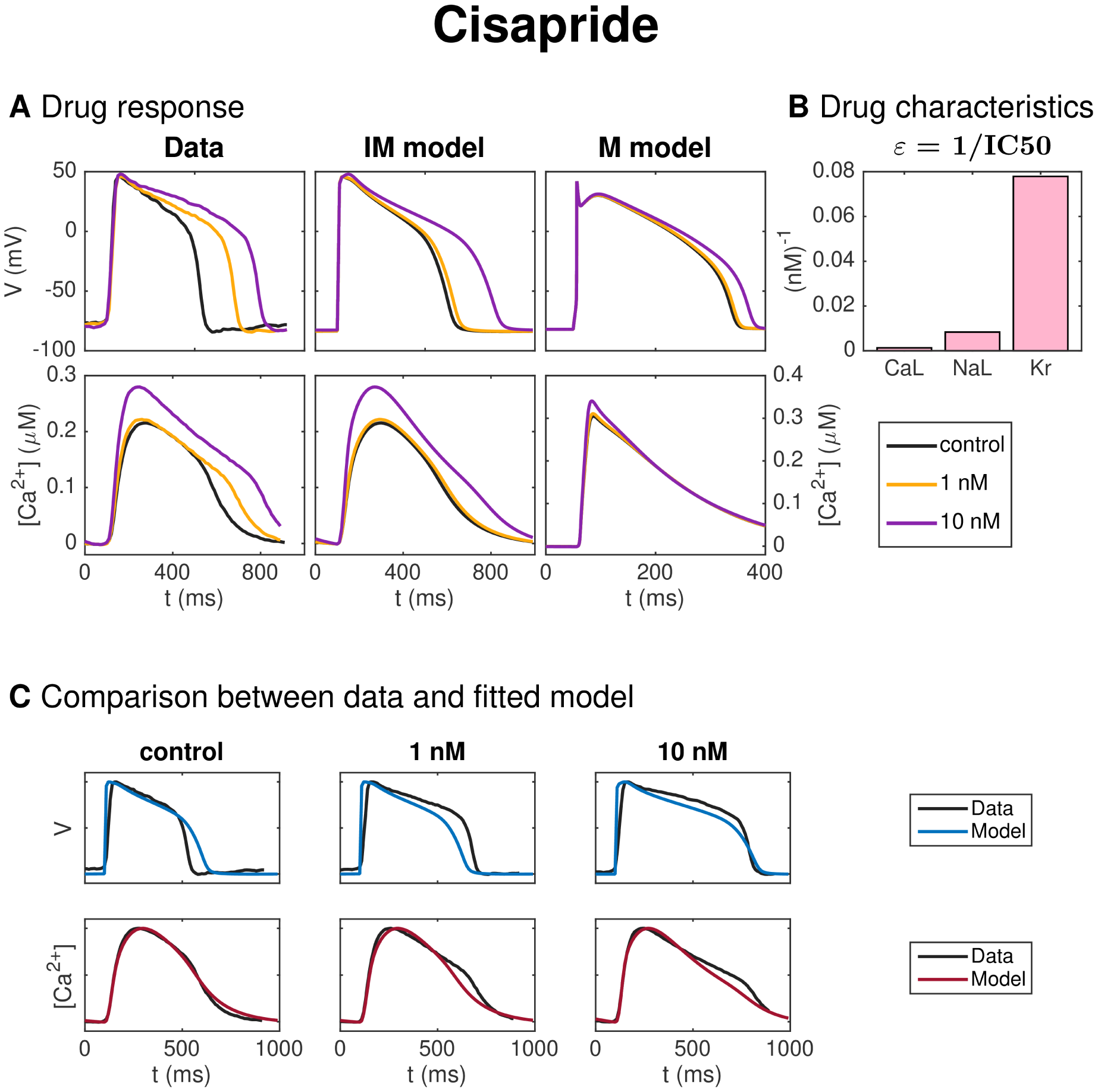
Identification and mapping of drug effects for the drug Cisapride based on optical measurements of the AP and Ca^2+^ transient of hiPSC-CMs following the same structure as Figure 7.

#### 3.3.4 Flecainide

Figure 10 shows the result for the inversion procedure applied to optical measurements of hiPSC-CMs exposed to the drug Flecainide. In panel A, we observe that the drug causes increased AP duration. In panel C, we observe that the fitted model fits the data quite well, excepting that the AP duration at high percentages of repolarization is longer for the data than for the model for the highest considered dose. In addition, the shape of the Ca^2+^ transient for the low doses is not entirely captured in the model. In panel B, we observe that the inversion procedure estimates that the drug primarily blocks *I*_Kr_ and, to some degree, *I*_CaL_.

**Figure 10:**
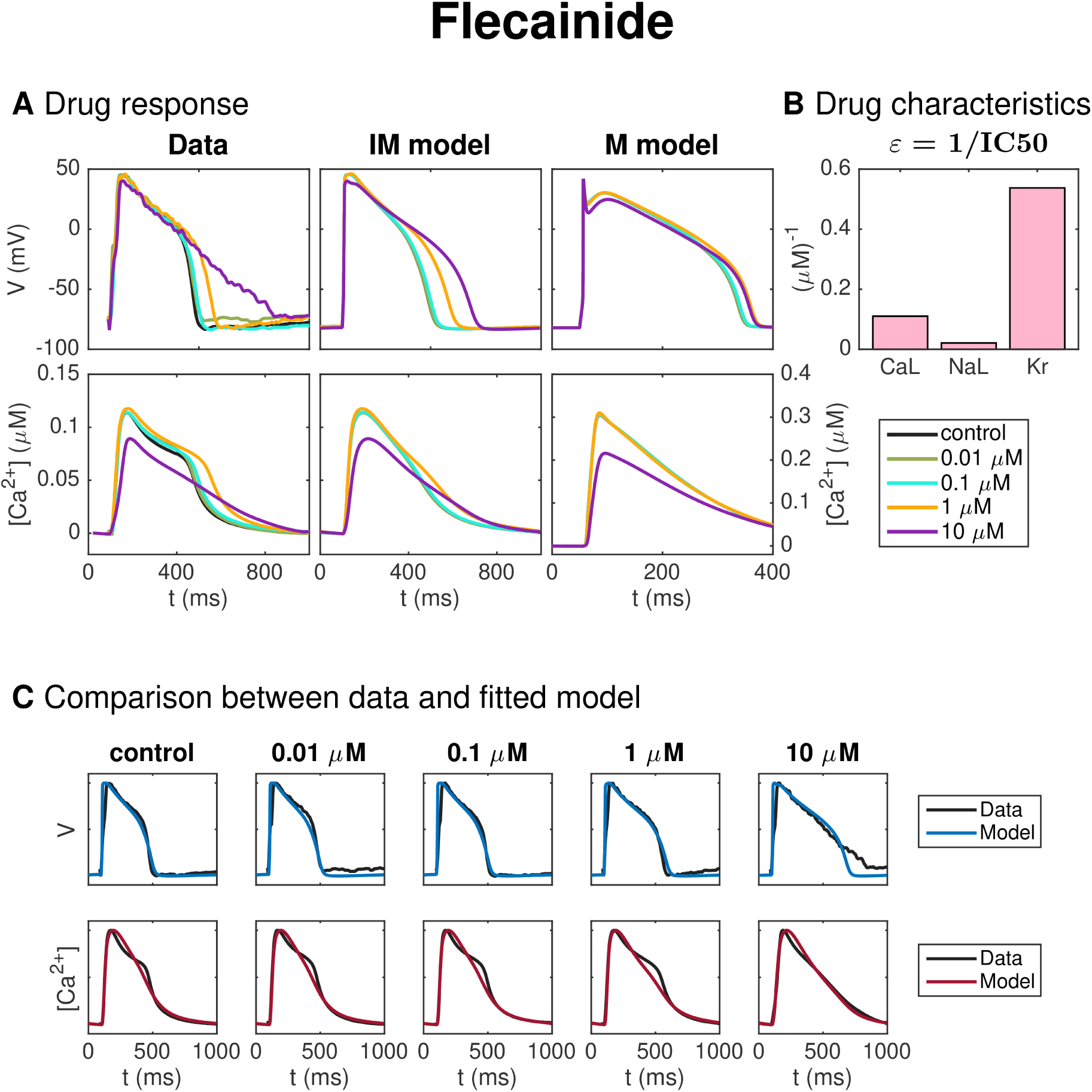
Identification and mapping of drug effects for the drug Flecainide based on optical measurements of the AP and Ca^2+^ transient of hiPSC-CMs following the same structure as Figure 7.

#### 3.3.5 Verapamil

Figure 11 shows the result of the inversion procedure applied to measurements of hiPSC-CMs exposed to the drug Verapamil. In panel A, we observe that the drug leads to decreased AP duration. Panel B shows that the inversion procedure predicts that Verapamil primarily blocks *I*_CaL_ and, to some extent, *I*_Kr_.

**Figure 11:**
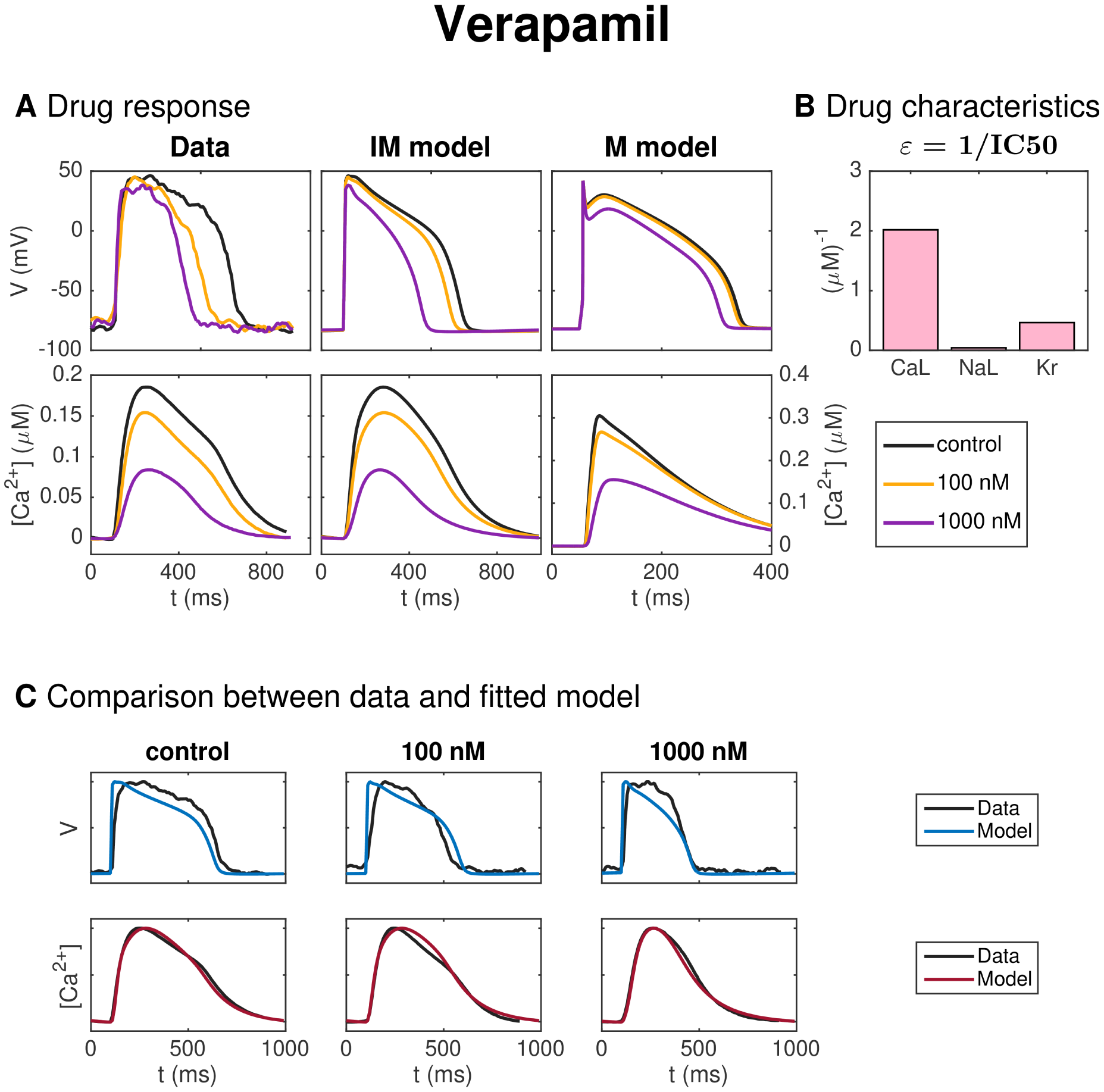
Identification and mapping of drug effects for the drug Verapamil based on optical measurements of the AP and Ca^2+^ transient of hiPSC-CMs, following the same structure as Figure 7.

### 3.4 Mapping of drug effects from hiPSC-CMs to mature cells

The rightmost plots of panel A of Figures 7–11 show the predicted drug effects for mature cells for each of the drugs considered. More specifically, the plots show the solution of the mature base model exposed to each drug’s effect (*ε*-values) as estimated by the inversion procedure for each of the drug doses included in the data set. To review, this represents the predicted drug response for a mature AP and Ca^2+^ transient exposed to each of the drugs, based on the optical measurements of the AP and Ca^2+^ transient as obtained in a microphysiological system of hiPSC-CMs. The predictions are made by first using the inversion procedure to estimate the effect of the drug on the *I*_CaL_, *I*_NaL_, and *I*_Kr_ currents in the immature case and then mapping the corresponding drug effects to a mature cell using the determined maturation map based on the assumptions of differences in the channel densities and geometry between immature and mature cells (see Section 2).

## 4 Discussion

Here, we have presented an improved version of the methods presented in [17] for estimating drug effects for adult human cardiomyocytes based on optical measurements of the AP and Ca^2+^ transient of immature cells (hiPSC-CMs in a microphysiological system). First, we introduce a new base model formulation for representing both mature and immature cells via different parameter sets. A model for intracellular Ca^2+^ dynamics is updated to a formulation constructed for stability with respect to parameter changes. In addition, we use IC50-based modeling of dose-dependent drug effects and find optimal parameters by running a coupled inversion of both the control data and the drug data for several different doses. The cost function measuring the difference between the data and the model has also been redefined, and we now apply a continuation-based minimization method to minimize the cost function.

### 4.1 Summary of method performance and main results: identification of drug effects based on simulated data and optical measurements of hiPSC-CMs

Figure 6 shows the result of the inversion procedure on simulated data. As noted above, we observe that the inversion procedure is able to identify the correct *ε*-values accurately, excepting the *ε*-value for Lidocaine, which is predicted to be considerably lower than the value used to generate the data. This suggests that it might be difficult to obtain correct values of *ε*_NaL_, as also supported by the low identifiability index for *I*_NaL_ reported in Figure 5. In addition, we observe that the inversion procedure predicts some block of *I*_NaL_ for the drug Cisapride, even though only *I*_Kr_ was blocked when the data was generated.

We additionally presented the use of the inversion procedure to identify drug effects on hiPSC-CMs from optical measurements of the AP and Ca^2+^ transients.

Figure 7 compares the experimentally measured data and the fitted model for each of the doses of Nifedipine applied to the microphysiological system. The model fits both the membrane potential and the Ca^2+^ data well for most doses applied, although the Ca^2+^ transient duration is longer in the model than in the data for the highest drug doses considered (see panel C). Furthermore, in panel B, we observe that the inversion procedure predicts that Nifedipine primarily blocks *I*_CaL_. In Table 3, we observe that the IC50 value for *I*_CaL_ is estimated to be 38 nM, in agreement with values found in literature (12 nM–60 nM [60, 61]). The IC50 value for *I*_NaL_ and *I*_Kr_ are estimated to be 23 600 nM and 40 200 nM, respectively — considerably larger than the doses considered in the data set. We have not found an IC50 value for *I*_NaL_ for comparison in literature, but the IC50 values found for *I*_Kr_ support the claim that the IC50 value is much larger than the drug doses included in the data set, although the literature values (275 000–440 000 nM [63, 60]) are higher than the value predicted by the inversion procedure.

Figure 8 shows similar results for inversion of measurements of hiPSC-CMs exposed to the drug Lidocaine. The AP duration is reduced by the drug and the inversion procedure predicts that the drug primarily blocks the *I*_NaL_ current. The IC50 value estimate for *I*_NaL_ is 4.3 *µ*M (see Table 3), in rough agreement with values found in literature (11 *µ*M [38]). We observe that the model fits the data quite well, but that the AP duration for the drug dose of 10 *µ*M is longer in the model than in the data.

Figure 9 shows the result of the inversion procedure applied to a data set for hiPSC-CMs exposed to the drug Cisapride. Considering the leftmost and center panels of Figure 9A, we observe that the prolongation of the AP duration is much more prominent in the data as compared to the fitted immature model for a drug dose of 1 nM. This is also confirmed in Figure 9C, where we observe that the model does not fit the membrane potential data for the control case and the 1 nM dose case well. The fit for the largest dose, however, is quite good. In Figure 9B, we observe that the inversion procedure predicts that Cisapride primarily blocks *I*_Kr_. In Table 3, we observe that the IC50 value for *I*_Kr_ is estimated to be 13 nM, in good agreement with values found in literature (6.5 nM–20 nM [64, 60, 38]).

Figure 10 shows the result for the method as applied to measurements of hiPSC-CMs exposed to the drug Flecainide, know to prolong the AP duration. In Table 3, we observe that the IC50 value for *I*_Kr_ predicted by the inversion procedure (1.9 *µ*M) is in quite good agreement with literature values (0.7-1.5 *µ*M [38, 60]), but that the predicted IC50 value for *I*_CaL_ (9 *µ*M) is too low compared to the reported literature values (26-27 *µ*M [38, 60]). In addition, the estimated IC50 value for *I*_NaL_ (47 *µ*M) is larger than the literature value of 19 *µ*M [38].

In Figure 11, the method is applied to measurements of hiPSC-CMs exposed to Verapamil. In panel A, the effect on the AP duration for the smallest dose (100 nM) appears to be more prominent in the data than in the fitted model. This is confirmed in panel C, where we observe that the fitted model seems to fit the Ca^2+^ data considerably better than the membrane potential data. In particular, the AP duration is too short in the control case and too long for the smallest dose of 100 nM. Panel B shows that the inversion procedure predicts that Verapamil primarily blocks *I*_CaL_ and to some extent *I*_Kr_. The predicted IC50 values from Table 3 (495 nM for *I*_CaL_ and 2150 nM for *I*_Kr_) are both higher than the corresponding values from literature (100– 202 nM for *I*_CaL_ [62, 60, 38] and 143–499 nM for *I*_Kr_ [65, 60, 38]).

The rightmost plots of panel A of Figures 7–11 show the predicted drug effects for mature cells for each of the well-characterized drugs considered. We observe that for some drugs (e.g., Nifedipine and Lidocaine), the drug effects for mature cells are predicted to be approximately as severe as for the immature cells. For other drugs, on the other hand, (e.g., Flecainide), the drug effect is predicted to be less severe for the mature cells than for immature cells, highlighting the critical importance of maturation phenotype in predictive biophysical models of hiPSC-CMs in pharmacological studies.

### 4.2 Modeling intracellular Ca^2+^ dynamics

The inversion algorithm requires thousands of simulations testing different parameters representing geometrical properties and channel densities, either in terms of membrane channels or in terms of channels or buffers involved in the intracellular Ca^2+^ machinery. In order for the inversion to work properly, it is essential that the AP model is stable with respect to variations in the parameters. In particular, it is important that the simulation does not fail because of instabilities in the model.

Modeling the intracellular Ca^2+^ dynamics of cardiac cells has been a long-standing challenge and a very active field of research for at least 40 years; for reviews see e.g., [57, 66, 67, 68, 69]. Ca^2+^ dynamics are a complex time-dependent, 3D and highly non-linear problem. Mathematical models have attempted to represent the dynamics using a system of ordinary differential equations. Essentially, the goal of these models has been to remove the spatial variance and compute solutions that are spatially averaged and therefore merely depend on time. The main motivation for this strategy is to achieve models that are practical to work with in terms of computational complexity. However, the strategy has run into serious modeling challenges that have subsequently been addressed with ingenuity in numerous models (see e.g., [70, 71, 72, 73, 74, 75, 42, 76, 57]). Also spatial models (see e.g., [77, 78, 79, 80]) and homogenized spatial models (see e.g., [81, 82, 83, 84, 85]) have been applied, and while these models clearly capture the intricate dynamics more convincingly, this comes at a computational cost that renders them impractical for the purpose of this study and many other applications, as tens of thousands of simulations with spatially resolved 3D models of the Ca^2+^ dynamics of cardiac cells is not computationally tractable at present. Below, we will discuss some important concepts involved in the intracellular Ca^2+^ dynamics of cardiac cells and some previously introduced modeling approaches for these dynamics.

#### 4.2.1 Ca^2+^ -induced Ca^2+^ release (CICR)

In the early phase of the upstroke of the AP, the membrane potential increases sufficiently for the voltage-sensitive dihydropyridine receptors (DHPR) to open the L-type Ca^2+^ channels on the membrane. Because of the huge gradient in the Ca^2+^ concentration between the intracellular and extracellular spaces, Ca^2+^ ions cross the membrane and flow into the cell. Inside the cell, the Ca^2+^ enters a tiny dyad (see Figure 2) located between the cell membrane and the sarcoplasmic reticulum (SR). Since the dyad is very small, the Ca^2+^ concentration increases rapidly and the increased concentration is sensed by the ryanodine receptors (RyRs) which in turn open and allow large amounts of Ca^2+^ to flow out of the SR. The increased Ca^2+^ concentration spreads by diffusion and recruits other RyRs to open and thus even more Ca^2+^ is poured into the bulk cytosolic space. This process is usually referred to as Ca^2+^ -induced Ca^2+^ release (CICR), and it takes place in many thousand local Ca^2+^ release units (CRUs) close to the cell membrane (see e.g., [86, 87, 56, 77]).

#### 4.2.2 High gain and graded release

The CICR is designed to provide both *high gain* and *graded release* (see e.g., [56, 57]). High gain means that, even when only a small amount of Ca^2+^ enters the cell through the cell membrane, this small amount leads to release of a much larger amount of Ca^2+^ from the SR. However, the release is also graded (see e.g., [88, 89, 86, 56, 57]) in the sense that the release of Ca^2+^ from the SR into the bulk cytosolic space depends continuously on the amount of Ca^2+^ flowing into the cell through the channels on the cell membrane. In other words, the amount of Ca^2+^ flowing into the cytosol during an AP is believed to be controlled by the flow through the membrane Ca^2+^ channels, even if most of the Ca^2+^ is released from the internal storage structures (the SR).

#### 4.2.3 Restoring the Ca^2+^ concentration

The AP is periodic and at the end of one cycle, all variables are brought back to the repolarized state of the cell. Ca^2+^ is pumped back to SR by the SERCA (sarcoplasmic reticulum Ca^2+^ ATPase) pump, and back to the extracellular space through the membrane Ca^2+^ pump and the Na^+^-Ca^2+^ exchanger.

#### 4.2.4 Common pool models

A standard approach to modeling CICR is illustrated in Figure 12. Here, the dynamics of the many CRUs are modeled by one representative unit, hence all CRUs are assumed to be in the same state. In the model, Ca^2+^ enters the dyad through L-type Ca^2+^ channels, which leads to an increased dyadic Ca^2+^ concentration, and thus the RyRs open and Ca^2+^ leaves the SR. Models of the form illustrated in Figure 12 is referred to as *common pool models* and are characterized by the fact that Ca^2+^ released through the RyRs (from the SR) enters the same, small, dyadic space that Ca^2+^ enters through the L-type Ca^2+^ channel. It has been known for a long time (see [70]) that it is impossible to obtain graded release using stable common pool models. The problem is that when the release of Ca^2+^ from the SR has started, the release from the SR will itself cause an increased dyadic Ca^2+^ concentration, and release will continue until some inactivation mechanism of the release (e.g., a sufficiently decreased SR Ca^2+^ concentration [40]) kicks in. Consequently, the release becomes an *all or nothing* process, depending only on whether the amount of Ca^2+^ entering the dyad through L-type Ca^2+^ channels is enough to trigger release. Therefore, graded release cannot be obtained using a model of the form given in Figure 12.

**Figure 12:**
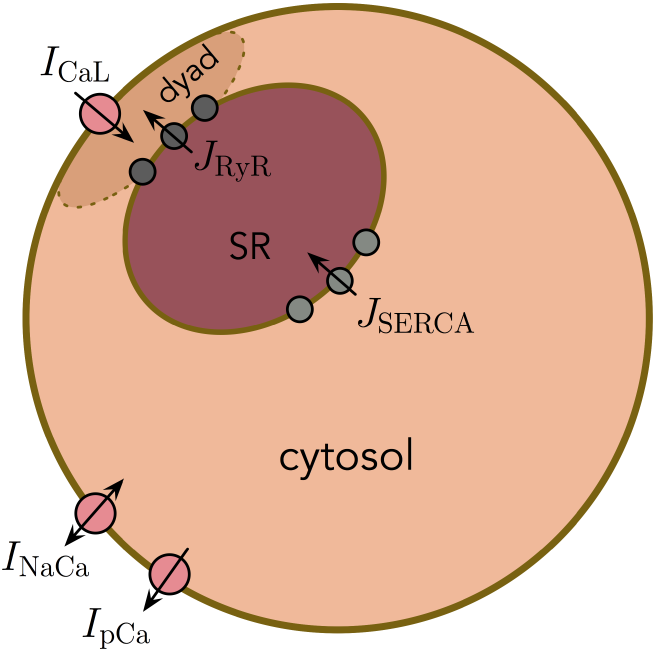
A prototypical sketch of a common pool model with Ca^2+^ flowing into the dyad through L-type Ca^2+^ channels; the RyR-channels are activated when the Ca^2+^ concentration increases, and Ca^2+^ is transported back to the SR from the cytosol via the SERCA pump and back to the extracellular space through the Na^+^-Ca^2+^ exchanger and the Ca^2+^ pump on the cell membrane.

#### 4.2.5 Local control models

The difficulties associated with the common pool models can be circumvented by allowing many CRUs in the model (see e.g., [77, 78, 56, 79]). By introducing a large number of CRUs that are weakly coupled and where the release mechanisms are governed by stochastic Markov models, it is possible to achieve both high gain and graded release. Suppose there are *∼*20.000 CRUs (as suggested in [79]) and every CRU has the elements illustrated in Figure 12 where the release mechanism of the L-type Ca^2+^ channel and the RyR are governed by Markov models. For simplicity we assume that the open probability of the L-type Ca^2+^ channels and the RyRs increases with increasing membrane potential and increasing dyadic Ca^2+^ concentration, respectively. Then, when the membrane potential increases slightly, the open probability of the L-type Ca^2+^ channels increases sufficiently for a few membrane channels to open, and thus Ca^2+^ will flow into the dyad of the associated CRUs. Locally, in these CRUs, the increased dyadic Ca^2+^ concentration will lead to increased open probability of the RyRs and when these channels open, the local SR of that particular CRU will be emptied. When the membrane potential increases more, the number of active CRUs will increase, and thus, the release will be graded by the membrane potential. So even if every single CRU is an all or nothing process, the integrated process is controlled by the membrane potential. Unfortunately, since these models requires a large number of CRUs, the computational cost of these models is prohibitive for our purposes.

#### 4.2.6 CICR in the base model

In the base model introduced above, we introduce two main modeling assumptions to obtain a model that exhibits both high gain and graded release without the high computational cost of local control models. First, the Ca^2+^ released from the SR is not released into the dyad, but is instead directed into a separate subsarcolemmal (SL) space. By directing the Ca^2+^ into this space, the Ca^2+^ entering the dyad though the membrane Ca^2+^ channels are clearly distinguishable from that released from the SR, and we avoid the graded-release problem associated with the common pool models. In addition, instead of inactivating the release from the SR by a decreased SR Ca^2+^ concentration, we introduce an assumption that each channel can only release a certain amount of Ca^2+^ during an AP cycle, introduced because the SR Ca^2+^ concentration can potentially vary significantly for the large parameter changes considered in the inversion procedure. In Figure 4, we observe that the model constructed from these assumptions exhibits both high gain and graded release for the immature and mature versions of the parameters.

### 4.3 Note on ongoing complementary studies

Other recent work has made strong progress in terms of enabling biophysical modeling approaches to assimilate and otherwise make best use of tailored experimental measurements of hiPSC-CMs. For example, in [90], the authors address the need to bridge the gap between the effect of drugs on human adult ventricular cardiomyocytes and the effect on animal or hiPSC experimental models often used in drug screening. This work also successfully generated accurate predictions of the effect of ion channel blocking drugs on human adult ventricular cardiomyocytes as based on simulations of hiPSC-CMs via a regression strategy.

Additional recent studies have advanced specific models and methodological approaches for hiPSC-CMs which incorporate experimental variability from multiple data sources (see, e.g., [91]) with the goal of identifying phenotypic mechanisms and identify key parameter sensitivity. The authors introduce a computational whole-cell electrophysiological model of hiPSC-CMs, composed of single exponential voltage-dependent gating variable rate constants, which are then parameterized to fit experimental measurements of hiPSC-CMs from multiple laboratories (and thus incorporate variability in the single-cell measurements of ionic currents observed experimentally). The authors compare immature and mature cell models to elucidate the primary properties underpinning the phenotype, a mechanistically central goal that was not an aim of our present study.

### 4.4 Limitations and notes on future work

CICR in the base model exhibits both high gain and graded release for the immature and mature parameter sets. However, these assumptions are introduced to obtain a stable model, and not necessarily to represent the underlying physiological mechanisms accurately. Future work necessitates assessment of the Ca^2+^ machinery in the base model and potential redevelopment to more accurately represent physiological Ca^2+^ release from the SR, as relevant.

In addition, we have only looked at a single dose escalation study for each of the drugs investigated. Future work will assess the variability of the inversion methodology in combination with noisy and incomplete experimental data obtained through these systems.

## Footnotes

1 Note that this does not apply to the regularization terms (87)–(88). These terms are assumed to be the same for all values of *θ*.

2 Note that this relies on either the flux balance term *H*_Ca,*b*_ being zero for the default base model or on the weight for this term being zero.

