## Supplementary material for "Improved computational identification of drug response using optical measurements of human stem cell derived cardiomyocytes in microphysiological systems"

### Supplementary information

#### 1 Base model formulation

In the following formulation of the base model, the membrane potential ($v$) is given in units of mV, and the $Ca^{2+}$concentrations are given in units of mM. All currents are expressed in units of A/F, and the $Ca^{2+}$fluxes are expressed as mmol/ms per total cell volume (i.e., in units of mM/ms). Time is given in ms. The parameters of the model are given in Tables 1–6.

##### 1.1 The membrane potential

The membrane potential is governed by the equation

$$\begin{aligned}\frac{dv}{dt} = -(&I_{\mathrm{Na}} + I_{\mathrm{NaL}} + I_{\mathrm{CaL}} + I_{\mathrm{to}} + I_{\mathrm{Kr}} + I_{\mathrm{Ks}} + I_{\mathrm{K1}} \\ &+ I_{\mathrm{NaCa}} + I_{\mathrm{NaK}} + I_{\mathrm{pCa}} + I_{\mathrm{bCl}} + I_{\mathrm{bCa}} + I_{\mathrm{f}} + I_{\mathrm{stim}}),\end{aligned} \tag{1}$$

where $I_{\mathrm{stim}}$ is an applied stimulus current, and $I_{\mathrm{Na}}$, $I_{\mathrm{NaL}}$, $I_{\mathrm{CaL}}$, $I_{\mathrm{to}}$, $I_{\mathrm{Kr}}$, $I_{\mathrm{Ks}}$, $I_{\mathrm{K1}}$, $I_{\mathrm{NaCa}}$, $I_{\mathrm{NaK}}$, $I_{\mathrm{pCa}}$, $I_{\mathrm{bCl}}$, $I_{\mathrm{bCa}}$, and $I_{\mathrm{f}}$ are membrane currents specified below. In our simulations, $I_{\mathrm{stim}}$ is given as a constant current of size $-40$ A/F for mature cells and $-5$ A/F for immature cells. The $I_{\mathrm{stim}}$ current is applied until the membrane potential reaches a value of $-40$ mV.

##### 1.2 Membrane currents

The currents through the voltage-gated ion channels on the cell membrane are in general given on the form

$$I = go(v - E),$$

where $g$ is the channel conductance, $v$ is the membrane potential and $E$ is the equilibrium potential of the channel. Furthermore, $o = \prod_i z_i$ is the open probability of the channels, where $z_i$ are gating variables, either given as a function of the membrane potential or governed by equations of the form

$$z_i' = \frac{1}{\tau_{z_i}}(z_{i,\infty} - z_i). \tag{2}$$

The parameters $\tau_{z_i}$ and $z_{i,\infty}$ will be specified for each of the gating variables of the model in Table 7.

**Fast sodium current** The formulation of the fast sodium current is an adjusted version of the model given in [1], supporting slower upstroke velocities more similar to those observed in the optical measurements of hiPSC-CMs. The current is given by

$$I_{\mathrm{Na}} = g_{\mathrm{Na}} o_{\mathrm{Na}} (v - E_{\mathrm{Na}}), \tag{3}$$

where the open probability is given by

$$o_{\mathrm{Na}} = m^3 j, \tag{4}$$

and $m$ and $j$ are gating variables governed by equations of the form (2).

**Late sodium current** The formulation of the late sodium current, $I_{\mathrm{NaL}}$, is based on [2] and is given by

$$I_{\mathrm{NaL}} = g_{\mathrm{NaL}} o_{\mathrm{NaL}} (v - E_{\mathrm{Na}}), \tag{5}$$

where the open probability is given by

$$o_{\mathrm{NaL}} = m_L h_L, \tag{6}$$

and $m_L$ and $h_L$ are gating variables governed by equations of the form (2).

**Transient outward potassium current** The formulation of the transient outward potassium current, $I_{\mathrm{to}}$, is based on [3] and is given by

$$I_{\mathrm{to}} = g_{\mathrm{to}} o_{\mathrm{to}} (v - E_{\mathrm{to}}), \tag{7}$$

where the open probability is given by

$$o_{\mathrm{to}} = q_{\mathrm{to}} r_{\mathrm{to}}, \tag{8}$$

and $q_{\mathrm{to}}$ and $r_{\mathrm{to}}$ are gating variables governed by equations of the form (2).

**Rapidly activating potassium current** The formulation of the rapidly activating potassium current, $I_{\mathrm{Kr}}$, is based on [3] and is given by

$$I_{\mathrm{Kr}} = g_{\mathrm{Kr}} o_{\mathrm{Kr}} (v - E_{\mathrm{K}}), \tag{9}$$

where

$$o_{\mathrm{Kr}} = x_{\mathrm{Kr1}} x_{\mathrm{Kr2}}, \tag{10}$$

and the dynamics of $x_{\mathrm{Kr1}}$ and $x_{\mathrm{Kr2}}$ are governed by equations of the form (2).

**Slowly activating potassium current** The formulation of the slowly activating potassium current, $I_{\mathrm{Ks}}$, is based on [1] and is given by

$$I_{\mathrm{Ks}} = g_{\mathrm{Ks}} o_{\mathrm{Ks}} (v - E_{\mathrm{Ks}}), \tag{11}$$

where

$$o_{\mathrm{Ks}} = x_{\mathrm{Ks}}^2, \tag{12}$$

and the dynamics of $x_{\mathrm{Ks}}$ is governed by an equation of the form (2).

**Inward rectifier potassium current** The formulation of the inward rectifier potassium current, $I_{\mathrm{K1}}$, is based on [1] and is given by

$$I_{\mathrm{K1}} = g_{\mathrm{K1}} o_{\mathrm{K1}} (v - E_{\mathrm{K}}), \tag{13}$$

where

$$o_{\mathrm{K1}} = \frac{a_{\mathrm{K1}}}{a_{\mathrm{K1}} + b_{\mathrm{K1}}}, \tag{14}$$

$$a_{\mathrm{K1}} = \frac{3.9}{1 + e^{0.6(v - E_K - 200)}}, \tag{15}$$

$$b_{\mathrm{K1}} = \frac{-1.5e^{0.0002(v - E_K + 100)} + e^{0.6(v - E_K - 10)}}{1 + e^{0.45(v - E_K)}}. \tag{16}$$

**Hyperpolarization activated funny current** The formulation for the hyperpolarization activated funny current, $I_{\mathrm{f}}$, is based on [3] and is given by

$$I_{\mathrm{f}} = g_{\mathrm{f}} o_{\mathrm{f}} (v - E_{\mathrm{f}}), \tag{17}$$

where

$$o_{\mathrm{f}} = x_{\mathrm{f}}, \tag{18}$$

and the dynamics of $x_{\mathrm{f}}$ is governed by an equation of the form (2).

**L-type $Ca^{2+}$current** The formualtion for the L-type $Ca^{2+}$current, $I_{\mathrm{CaL}}$, is based on the formulation in [1] and is given by

$$I_{\mathrm{CaL}} = g_{\mathrm{CaL}} o_{\mathrm{CaL}} \frac{(2F)^2 v}{RT} \frac{0.341 c_d e^{\frac{2Fv}{RT}} - 0.341 c_e}{e^{\frac{2Fv}{RT}} - 1}, \tag{19}$$

where

$$o_{\mathrm{CaL}} = df(1 - f_{\mathrm{Ca}}), \tag{20}$$

and the dynamics of $d$, $f$ and $f_{\mathrm{Ca}}$ are governed by equations of the form (2).

**Background currents** The formulation of the background currents, $I_{\mathrm{bCa}}$ and $I_{\mathrm{bCl}}$, are based on [1] and are given by

$$I_{\mathrm{bCa}} = g_{\mathrm{bCa}}(v - E_{\mathrm{Ca}}), \tag{21}$$

$$I_{\mathrm{bCl}} = g_{\mathrm{bCl}}(v - E_{\mathrm{Cl}}). \tag{22}$$

**Sodium-calcium exchanger** The formulation of the $Na^+$-$Ca^{2+}$ exchanger current, $I_{\mathrm{NaCa}}$, is based on [1] and is given by

$$I_{\mathrm{NaCa}} = \bar{I}_{\mathrm{NaCa}} \frac{e^{\frac{\nu F v}{RT}} [\mathrm{Na}^+]_i^3 c_e - e^{\frac{(\nu-1)Fv}{RT}} [\mathrm{Na}^+]_e^3 c_{sl}}{s_{\mathrm{NaCa}} \left(1 + \left(\frac{K_{\mathrm{act}}}{c_{sl}}\right)^2\right) \left(1 + k_{\mathrm{sat}} e^{\frac{(\nu-1)Fv}{RT}}\right)}, \tag{23}$$

where

$$s_{\mathrm{NaCa}} = K_{\mathrm{Ca},i} [\mathrm{Na}^+]_e^3 \left(1 + \left(\frac{[\mathrm{Na}^+]_i}{K_{\mathrm{Na},i}}\right)^3\right) + K_{\mathrm{Na},e}^3 c_{sl} \left(1 + \frac{c_{sl}}{K_{\mathrm{Ca},i}}\right)$$
$$+ K_{\mathrm{Ca},e} [\mathrm{Na}^+]_i^3 + [\mathrm{Na}^+]_i^3 c_e + [\mathrm{Na}^+]_e^3 c_{sl}.$$

**Sarcolemmal $Ca^{2+}$pump** The formulation of the current through the sarcolemmal $Ca^{2+}$pump, $I_{\mathrm{pCa}}$, is based on [1] and is given by

$$I_{\mathrm{pCa}} = \bar{I}_{\mathrm{pCa}} \frac{c_{sl}^2}{K_{\mathrm{pCa}}^2 + c_{sl}^2}. \tag{24}$$

**Sodium-potassium pump** The current through the $Na^+$-$K^+$ pump, $I_{\mathrm{NaK}}$, is based on [1] and is given by

$$I_{\mathrm{NaK}} = \bar{I}_{\mathrm{NaK}} \frac{f_{\mathrm{NaK}}}{1 + \left(\frac{K_{\mathrm{Na,i}}^{\mathrm{NaK}}}{[\mathrm{Na}^+]_\mathrm{i}}\right)^4} \frac{[\mathrm{K}^+]_\mathrm{e}}{[\mathrm{K}^+]_\mathrm{e} + K_{\mathrm{K,e}}}, \tag{25}$$

where

$$f_{\mathrm{NaK}} = \frac{1}{1 + 0.12 e^{-0.1 \frac{Fv}{RT}}} + \frac{0.037}{7} \left(e^{\frac{[\mathrm{Na}^+]_\mathrm{e}}{67}} - 1\right) e^{-\frac{Fv}{RT}}. \tag{26}$$

#### 1.3 Ca$^{2+}$dynamics

The Ca$^{2+}$dynamics are governed by

$$\frac{dc_d}{dt} = \frac{1}{V_d}(J_{\text{CaL}} - J_d^b - J_d^c), \qquad \frac{db_d}{dt} = \frac{1}{V_d}J_d^b, \tag{27}$$

$$\frac{dc_{sl}}{dt} = \frac{1}{V_{sl}}(J_e^{sl} - J_{sl}^c - J_{sl}^b + J_s^{sl}), \qquad \frac{db_{sl}}{dt} = \frac{1}{V_{sl}}J_{sl}^b, \tag{28}$$

$$\frac{dc_c}{dt} = \frac{1}{V_c}(J_{sl}^c + J_d^c - J_c^n - J_c^b), \qquad \frac{db_c}{dt} = \frac{1}{V_c}J_c^b, \tag{29}$$

$$\frac{dc_s}{dt} = \frac{1}{V_s}(J_n^s - J_s^{sl} - J_s^b), \qquad \frac{db_s}{dt} = \frac{1}{V_s}J_s^b, \tag{30}$$

$$\frac{dc_n}{dt} = \frac{1}{V_n}(J_c^n - J_n^s), \tag{31}$$

where $c_d$ is the concentration of free Ca$^{2+}$in the dyad, $b_d$ is the concentration of Ca$^{2+}$bound to a buffer in the dyad, $c_{sl}$ is the concentration of free Ca$^{2+}$in the SL compartment, $b_{sl}$ is the concentration of Ca$^{2+}$bound to a buffer in the SL compartment, $c_c$ is the concentration of free Ca$^{2+}$in the bulk cytosol, $b_c$ is the concentration of Ca$^{2+}$bound to a buffer in the bulk cytosol, $c_s$ is the concentration of free Ca$^{2+}$in the jSR, $b_s$ is the concentration of Ca$^{2+}$bound to a buffer in the jSR, and $c_n$ is the concentration of free Ca$^{2+}$in the nSR. The expressions for the fluxes are specified below.

#### 1.4 Ca$^{2+}$fluxes

**Flux through the SERCA pumps** The flux from the bulk cytosol to the nSR through the SERCA pumps is given by

$$J_c^n = \bar{J}_{\text{SERCA}} \frac{\left(\frac{c_c}{K_c}\right)^2 - \left(\frac{c_n}{K_n}\right)^2}{1 + \left(\frac{c_c}{K_c}\right)^2 + \left(\frac{c_n}{K_n}\right)^2}. \tag{32}$$

**Flux through the RyRs** The flux from the jSR to the SL compartment is given by

$$J_s^{sl} = J_{\text{RyR}} + J_{\text{leak}}, \tag{33}$$

where $J_{\text{RyR}}$ represents the flux through the active RyR channels and $J_{\text{leak}}$ represents the flux through the RyR channels that are always open, given by

$$J_{\text{RyR}} = p \cdot r \cdot \alpha_{\text{RyR}}(c_s - c_{sl}), \tag{34}$$

$$J_{\text{leak}} = \gamma_{\text{RyR}} \cdot \alpha_{\text{RyR}}(c_s - c_{sl}), \tag{35}$$

respectively. Here, $p$ is the open probability of the active RyR channels given by

$$p = \frac{c_d^3}{c_d^3 + \kappa_{\text{RyR}}^3}, \tag{36}$$

and $r$ represents the fraction of RyR channels that are not inactivated and is governed by the equation

$$\frac{dr}{dt} = -\frac{J_{\text{RyR}}}{\beta_{\text{RyR}}} + \frac{\eta_{\text{RyR}}}{p}(1-r). \tag{37}$$

**Passive diffusion fluxes between compartments** The passive diffusion fluxes between compartments are given by

$$J_d^c = \alpha_d^c(c_d - c_c), \tag{38}$$
$$J_{sl}^c = \alpha_{sl}^c(c_{sl} - c_c), \tag{39}$$
$$J_n^s = \alpha_n^s(c_n - c_s). \tag{40}$$

**Buffer fluxes** The fluxes of free $Ca^{2+}$ binding to a $Ca^{2+}$buffer are given by

$$J_d^b = V_d(k_{\text{on}}^d c_d(B_{\text{tot}}^d - b_d) - k_{\text{off}}^d b_d), \tag{41}$$
$$J_{sl}^b = V_{sl}(k_{\text{on}}^{sl} c_{sl}(B_{\text{tot}}^{sl} - b_{sl}) - k_{\text{off}}^{sl} b_{sl}), \tag{42}$$
$$J_c^b = V_c(k_{\text{on}}^c c_c(B_{\text{tot}}^c - b_c) - k_{\text{off}}^c b_c), \tag{43}$$
$$J_s^b = V_s(k_{\text{on}}^s c_s(B_{\text{tot}}^s - b_s) - k_{\text{off}}^s b_s). \tag{44}$$

**Membrane fluxes** The membrane fluxes, $J_{\text{CaL}}$, $J_{\text{bCa}}$, $J_{\text{pCa}}$, and $J_{\text{NaCa}}$, are given by

$$J_{\text{CaL}} = -\frac{\chi C_m}{2F} I_{\text{CaL}}, \qquad J_{\text{pCa}} = -\frac{\chi C_m}{2F} I_{\text{pCa}}, \tag{45}$$
$$J_{\text{bCa}} = -\frac{\chi C_m}{2F} I_{\text{bCa}}, \qquad J_{\text{NaCa}} = \frac{\chi C_m}{F} I_{\text{NaCa}}, \tag{46}$$

where $I_{\text{CaL}}$, $I_{\text{bCa}}$, $I_{\text{pCa}}$, and $I_{\text{NaCa}}$ are defined by the expressions given above. Furthermore,

$$J_e^{sl} = J_{\text{NaCa}} + J_{\text{pCa}} + J_{\text{bCa}}. \tag{47}$$

| Parameter | Description | Value |
| --- | --- | --- |
| $V_d$ | Volume fraction of the dyadic subspace | 0.001 |
| $V_{sl}$ | Volume fraction of the SL compartment | 0.028 |
| $V_c$ | Volume fraction of the bulk cytosol | 0.917 |
| $V_s$ | Volume fraction of the jSR | 0.004 |
| $V_n$ | Volume fraction of the nSR | 0.05 |
| $\chi$ | Cell surface to volume ratio | 0.6 $\mu$m$^{-1}$ |

Table S1: Default geometry parameters of the base model.

#### 1.5 Nernst equilibrium potentials

The Nernst equilibrium potentials for the ion channels are defined as

$$E_{\mathrm{Na}} = \frac{RT}{F}\log\left(\frac{[\mathrm{Na}^+]_\mathrm{e}}{[\mathrm{Na}^+]_\mathrm{i}}\right), \tag{48}$$

$$E_{\mathrm{Ca}} = \frac{RT}{2F}\log\left(\frac{[\mathrm{Ca}^{2+}]_\mathrm{e}}{c_{sl}}\right), \tag{49}$$

$$E_{\mathrm{K}} = \frac{RT}{F}\log\left(\frac{[\mathrm{K}^+]_\mathrm{e}}{[\mathrm{K}^+]_\mathrm{i}}\right), \tag{50}$$

$$E_{\mathrm{Ks}} = \frac{RT}{F}\log\left(\frac{[\mathrm{K}^+]_\mathrm{e} + 0.018[\mathrm{Na}^+]_\mathrm{e}}{[\mathrm{K}^+]_\mathrm{i} + 0.018[\mathrm{Na}^+]_\mathrm{i}}\right), \tag{51}$$

$$E_{\mathrm{Cl}} = \frac{RT}{F}\log\left(\frac{[\mathrm{Cl}^+]_\mathrm{e}}{[\mathrm{Cl}^+]_\mathrm{i}}\right), \tag{52}$$

$$E_{\mathrm{f}} = -17 \text{ mV}, \tag{53}$$

for the parameter values given in Table 2.

| Parameter | Description | Value |
|---|---|---|
| $C_m$ | Specific membrane capacitance | 0.01 $\mu$F/$\mu$m$^2$ |
| $F$ | Faraday's constant | 96.485 C/mmol |
| $R$ | Universal gas constant | 8.314 J/(mol·K) |
| $T$ | Temperature | 310 K |
| $[Ca^{2+}]_e$ | Extracellular $Ca^{2+}$concentration | 1.8 mM |
| $[Na^{+}]_e$ | Extracellular sodium concentration | 140 mM |
| $[Na^{+}]_i$ | Intracellular sodium concentration | 8 mM |
| $[K^{+}]_e$ | Extracellular potassium concentration | 5.4 mM |
| $[K^{+}]_i$ | Intracellular potassium concentration | 120 mM |
| $[Cl^{-}]_e$ | Extracellular chloride concentration | 150 mM |
| $[Cl^{-}]_i$ | Intracellular chloride concentration | 15 mM |

Table S2: Physical constants and ionic concentrations of the base model.

| Parameter | Value | Parameter | Value |
|---|---|---|---|
| $g_{\mathrm{Na}}$ | 12.6 mS/$\mu$F | $g_{\mathrm{CaL}}$ | 0.12 nL/($\mu$F ms) |
| $g_{\mathrm{NaL}}$ | 0.025 mS/$\mu$F | $g_{\mathrm{bCa}}$ | 0.00055 mS/$\mu$F |
| $g_{\mathrm{to}}$ | 0.27 mS/$\mu$F | $\bar{I}_{\mathrm{NaCa}}$ | 4.9 $\mu$A/$\mu$F |
| $g_{\mathrm{Kr}}$ | 0.025 mS/$\mu$F | $\bar{I}_{\mathrm{pCa}}$ | 0.068 $\mu$A/$\mu$F |
| $g_{\mathrm{Ks}}$ | 0.003 mS/$\mu$F | $\bar{J}_{\mathrm{SERCA}}$ | 0.00024 mM/ms |
| $g_{\mathrm{K1}}$ | 0.37 mS/$\mu$F | $\alpha_{\mathrm{RyR}}$ | 0.0075 ms$^{-1}$ |
| $g_{\mathrm{f}}$ | 0.0001 mS/$\mu$F | $\alpha_d^c$ | 0.0017 ms$^{-1}$ |
| $g_{\mathrm{bCl}}$ | 0.007 mS/$\mu$F | $\alpha_{sl}^c$ | 0.15 ms$^{-1}$ |
| $\bar{I}_{\mathrm{NaK}}$ | 1.8 $\mu$A/$\mu$F | $\alpha_n^s$ | 0.012 ms$^{-1}$ |

Table S3: Conductance values and similar parameters for each of the membrane currents and intracellular $Ca^{2+}$fluxes of the base model.

| Parameter | Flux | Value |
|---|---|---|
| $K_c$ | $J_c^n$ | 0.00025 mM |
| $K_n$ | $J_c^n$ | 1.7 mM |
| $\beta_{\mathrm{RyR}}$ | $J_s^{sl}$ | 0.038 mM |
| $\gamma_{\mathrm{RyR}}$ | $J_s^{sl}$ | 0.001 |
| $\kappa_{\mathrm{RyR}}$ | $J_{\mathrm{RyR}}$ | 0.015 mM |
| $\eta_{\mathrm{RyR}}$ | $J_s^{sl}$ | 0.00001 $\mathrm{ms}^{-1}$ |

Table S4: Parameters for the intracellular $Ca^{2+}$fluxes of the base model.

| Parameter | Current | Value |
|---|---|---|
| $k_{\mathrm{sat}}$ | $I_{\mathrm{NaCa}}$ | 0.3 |
| $\nu$ | $I_{\mathrm{NaCa}}$ | 0.3 |
| $K_{\mathrm{act}}$ | $I_{\mathrm{NaCa}}$ | 0.00015 mM |
| $K_{\mathrm{Ca},i}$ | $I_{\mathrm{NaCa}}$ | 0.0036 mM |
| $K_{\mathrm{Ca},e}$ | $I_{\mathrm{NaCa}}$ | 1.3 mM |
| $K_{\mathrm{Na},i}$ | $I_{\mathrm{NaCa}}$ | 12.3 mM |
| $K_{\mathrm{Na},e}$ | $I_{\mathrm{NaCa}}$ | 87.5 mM |
| $K_{\mathrm{Na},i}^{\mathrm{NaK}}$ | $I_{\mathrm{NaK}}$ | 11 mM |
| $K_{\mathrm{K},e}$ | $I_{\mathrm{NaK}}$ | 1.5 mM |
| $K_{\mathrm{pCa}}$ | $I_{\mathrm{pCa}}$ | 0.0005 mM |

Table S5: Parameters for the membrane currents of the base model.

| Parameter | Compartment | Value |
|---|---|---|
| $B^{c}_{\text{tot}}$ | Bulk cytosol | 0.07 mM |
| $k^{c}_{\text{on}}$ | Bulk cytosol | 40 $\text{ms}^{-1}\text{mM}^{-1}$ |
| $k^{c}_{\text{off}}$ | Bulk cytosol | 0.03 $\text{ms}^{-1}$ |
| $B^{d}_{\text{tot}}$ | Dyad | 1.2 mM |
| $k^{d}_{\text{on}}$ | Dyad | 100 $\text{ms}^{-1}\text{mM}^{-1}$ |
| $k^{d}_{\text{off}}$ | Dyad | 1 $\text{ms}^{-1}$ |
| $B^{sl}_{\text{tot}}$ | Subsarcolemmal space | 0.9 mM |
| $k^{sl}_{\text{on}}$ | Subsarcolemmal space | 100 $\text{ms}^{-1}\text{mM}^{-1}$ |
| $k^{sl}_{\text{off}}$ | Subsarcolemmal space | 0.15 $\text{ms}^{-1}$ |
| $B^{s}_{\text{tot}}$ | Junctional SR | 27 mM |
| $k^{s}_{\text{on}}$ | Junctional SR | 100 $\text{ms}^{-1}\text{mM}^{-1}$ |
| $k^{s}_{\text{off}}$ | Junctional SR | 65 $\text{ms}^{-1}$ |

Table S6: Parameters for the $Ca^{2+}$buffers of the base model.

| Current | Gate | $z_\infty$ | $\alpha_z$ | $\beta_z$ | $\tau_z$ |
|---|---|---|---|---|---|
| $I_{\text{Na}}$ | $m$ | $\frac{1}{\left(1+e^{(-57-v)/9}\right)^2}$ | $0.13e^{-((v+46)/16)^2}$ | $0.06e^{-((v-5)/51)^2}$ | $\alpha_m+\beta_m$ |
| | $j$ | $\frac{1}{\left(1+e^{(v+72)/7}\right)^2}$ | $0$, if $v \geq -40$<br>$\frac{\begin{pmatrix} -2.5\cdot 10^4 e^{0.2v} \\ -7\cdot 10^{-6}e^{-0.04v} \end{pmatrix}(v+38)}{1+e^{0.3(v+79)}}$, otherwise | $\frac{0.6e^{0.06v}}{1+e^{-0.1(v+32)}}$, if $v \geq -40$<br>$\frac{0.02e^{-0.01v}}{1+e^{-0.14(v+40)}}$, otherwise | $\frac{1}{\alpha_j+\beta_j}$ |
| $I_{\text{NaL}}$ | $m_L$ | $\frac{1}{1+e^{(-43-v)/5}}$ | $\frac{1}{6.8e^{(v+12)/35}}$ | $8.6e^{-(v+77)/6}$ | $\alpha_m+\beta_m$ |
| | $h_L$ | $\frac{1}{1+e^{(v+88)/7.5}}$ | | | 200 ms |
| $I_{\text{CaL}}$ | $d$ | $\frac{1}{1+e^{-(v+5)/6}}$ | $\frac{1-e^{-\frac{v+5}{6}}}{0.035(v+5)}$ | | $\alpha_d$ |
| | $f$ | $\frac{1}{1+e^{(v+35)/9}}+\frac{0.6}{1+e^{(50-v)/20}}$ | $\frac{1}{0.02e^{-(0.034(v+14.5)^2)}+0.02}$ | | $\alpha_f$ |
| | $f_{\text{Ca}}$ | $\frac{1.7c_d}{1.7c_d+0.012}$ | $\frac{1}{1.7c_d+0.012}$ | | $\alpha_{\text{Ca}}$ |
| $I_{\text{to}}$ | $q_{\text{to}}$ | $\frac{1}{1+e^{(v+53)/13}}$ | $\frac{39}{0.57e^{-0.08(v+44)}+0.065e^{0.1(v+46)}}$ | 6 | $\alpha_{q_{\text{to}}}+\beta_{q_{\text{to}}}$ |
| | $r_{\text{to}}$ | $\frac{1}{1+e^{-(v-22.3)/18.75}}$ | $\frac{14.4}{e^{0.09(v+30.61)}+0.37e^{-0.12(v+24)}}$ | 2.75 | $\alpha_{r_{\text{to}}}+\beta_{r_{\text{to}}}$ |
| $I_{\text{Kr}}$ | $x_{\text{Kr1}}$ | $\frac{1}{1+e^{-(v+20.7)/4.9}}$ | $\frac{450}{1+e^{-(v+45)/10}}$ | $\frac{6}{1+e^{(v+30)/11.5}}$ | $\alpha_{x_{\text{Kr1}}}\cdot\beta_{x_{\text{Kr1}}}$ |
| | $x_{\text{Kr2}}$ | $\frac{1}{1+e^{(v+88)/50}}$ | $\frac{3}{1+e^{-(v+60)/20}}$ | $\frac{1.12}{1+e^{(v-60)/20}}$ | $\alpha_{x_{\text{Kr2}}}\cdot\beta_{x_{\text{Kr2}}}$ |
| $I_{\text{Ks}}$ | $x_{\text{Ks}}$ | $\frac{1}{1+e^{-(v+3.8)/14}}$ | $\frac{990}{1+e^{-(v+2.4)/14}}$ | | $\alpha_{x_{\text{Ks}}}$ |
| $I_{\text{f}}$ | $x_{\text{f}}$ | $\frac{1}{1+e^{(v+78)/5}}$ | $\frac{1900}{1+e^{(v+15)/10}}$ | | $\alpha_{x_{\text{Ks}}}$ |

Table S7: Specification of the parameters $z_\infty$ and $\tau_z$, for $z = m$, $j$, $m_L$, $h_L$, $d$, $f$, $f_{\text{Ca}}$, $q_{\text{to}}$, $r_{\text{to}}$, $x_{\text{Kr1}}$, $x_{\text{Kr2}}$, $x_{\text{Ks}}$ and $x_{\text{f}}$ in the equations for the gating variables (2).

#### 2 Supplementary figures

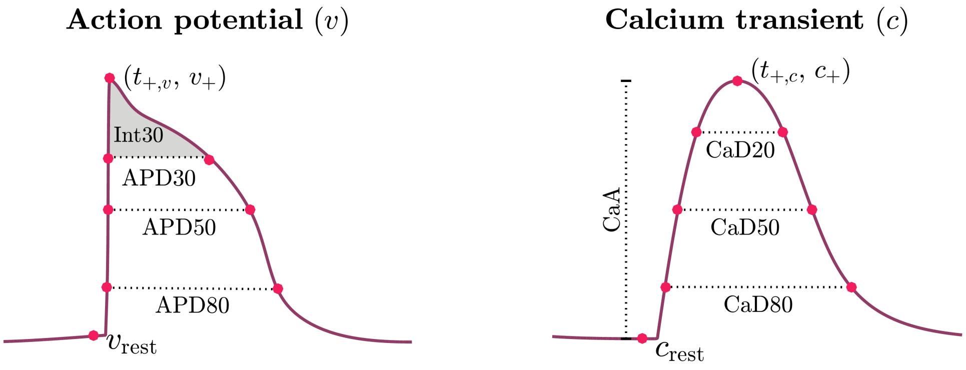


Figure S1: Illustration of some of the quantities used to define the terms of the cost function from the AP and $Ca^{2+}$transient.

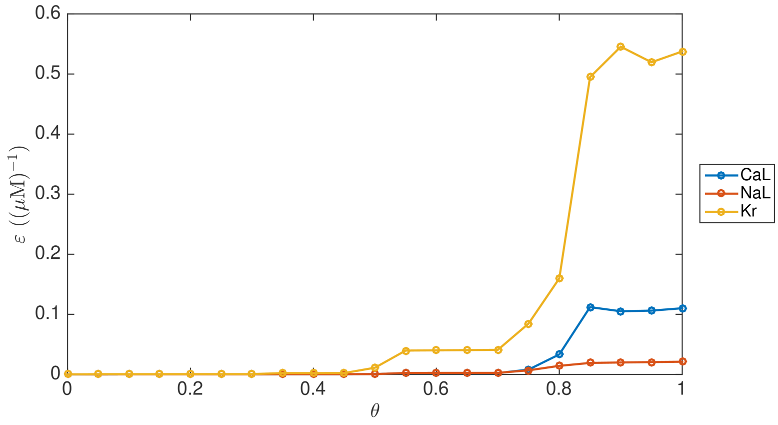


Figure S2: Example of how the optimal $\varepsilon$-values gradually move from zero to the optimal values for the data as $\theta$ is increased from zero to one in the continuation-based inversion procedure.
